# Acute social isolation evokes midbrain craving responses similar to hunger

**DOI:** 10.1101/2020.03.25.006643

**Authors:** Livia Tomova, Kimberly L. Wang, Todd Thompson, Gillian A. Matthews, Atsushi Takahashi, Kay M. Tye, Rebecca Saxe

**Author notes:** Correspondence to: Livia Tomova.

## Abstract

When people are forced to be isolated from one another, do they crave social interactions? To address this question, we used functional magnetic resonance imaging (fMRI) to measure neural responses evoked by food and social cues after participants (n=40) experienced ten hours of mandated fasting or total social isolation. After isolation, people felt lonely and craved social interaction. Midbrain regions showed selective activation to food cues after fasting and to social cues after isolation; these responses were correlated with self-reported craving. By contrast, striatal and cortical regions differentiated between craving food versus social interaction. Across deprivation sessions, we find that deprivation narrows and focuses the brain’s motivational responses to the deprived target. Our results support the intuitive idea that acute isolation causes social craving, similar to the way fasting causes hunger.

---

How are people affected by a period of forced social isolation? Chronic social isolation and loneliness are associated with lower physical^1,2^ and mental^2–4^ health, but little is known about the consequences of acute mandatory isolation. Positive social interactions in and of themselves may be basic human needs, analogous to other basic needs like food consumption or sleep^5,6^. If so, the absence of positive social interaction may create a want, or “craving”, that motivates behavior to repair what is lacking^5^. Cues associated with positive social interaction (e.g., smiling faces) activate neural reward systems^7, for review^. However, research on the neural representation of *unmet* human social needs is scarce^8^.

In social animals, social interactions act as primary rewards^9–11^: they are inherently pleasurable and motivate behavior in the absence of any other reward. Extended periods of isolation, especially during development, can dramatically disrupt behavior and brain function^12–14^. Even a brief acute period of social isolation causes an aversive, ‘loneliness-like’ brain state in adult mice, causing the mice to seek social interaction^15^ which is mediated specifically by dopaminergic midbrain neurons^16^, similar to other kinds of craving^17, for review^.

However, the homology to human loneliness has been disputed^8^, and it is not possible to assess whether a mouse subjectively feels lonely when isolated. Would acute isolation evoke a similar response in humans? The intuitive idea that depriving social needs evokes social craving, analogous to the way fasting evokes food craving and is mediated by similar dopaminergic midbrain regions, has never been directly tested in humans. The few prior studies investigating the correlation between self-reported chronic loneliness and brain responses to social stimuli yield contradictory findings^18–20^ and are limited by the ambiguity of observed correlations: if brain responses do differ, are the differences antecedents or effects of loneliness?

To address these questions, we experimentally induced social isolation experimentally in a within-subject design. 40 healthy socially-connected young adults (ages 18-40, 27 women) underwent 10 hours of social isolation and an fMRI scan with a cue-induced craving paradigm. Each participant also underwent 10 hours of food fasting and a subsequent MRI scan. Figure 1 shows an overview of the experimental procedures (see Methods for details). All predictions and methods were preregistered on the Open Science Framework (https://osf.io/cwg9e/).

**Figure 1.**
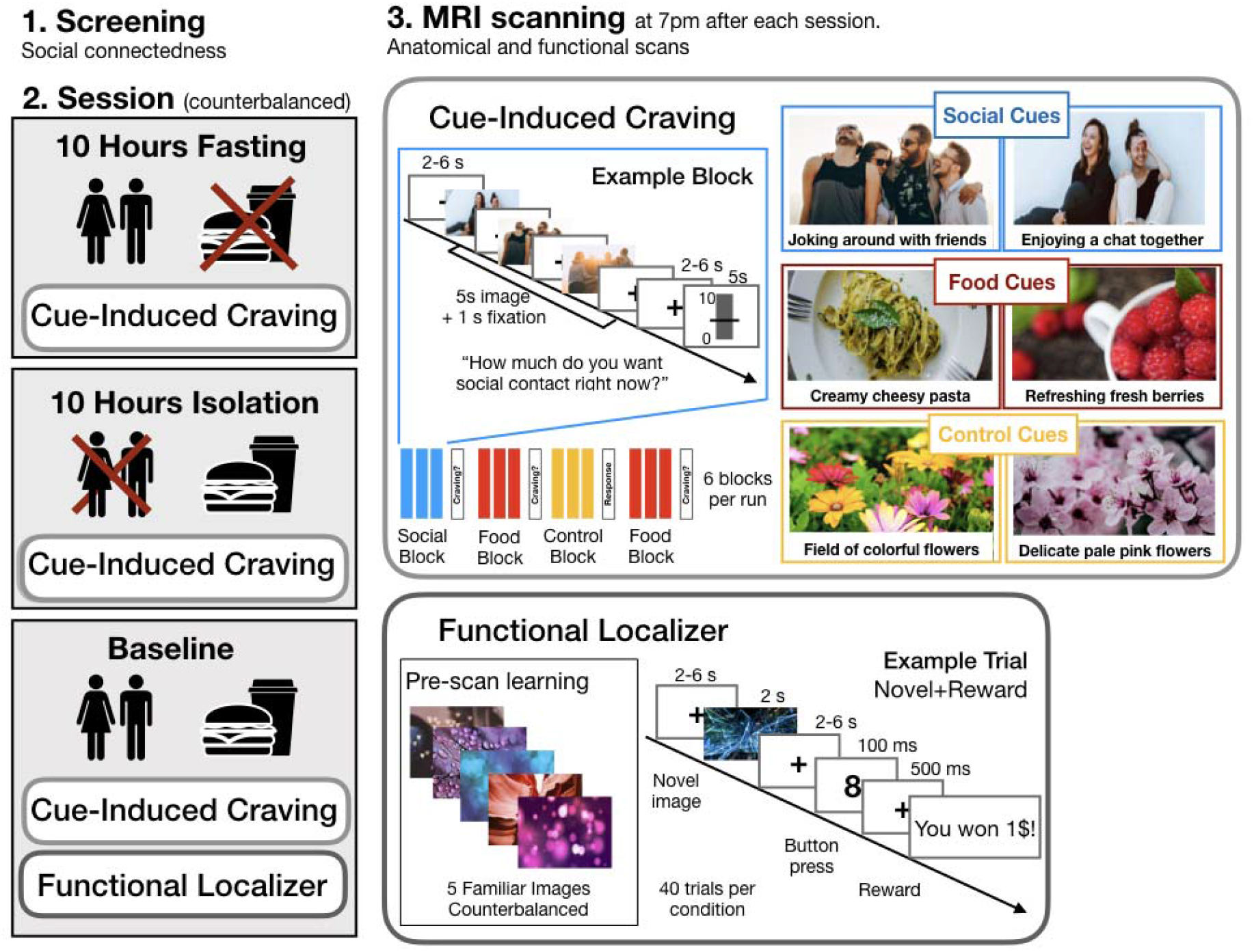
Overview of the experimental procedures. First, individuals underwent a screening for their social connectedness (measured by social network size and self-reported loneliness; see Methods for details). Each participant (n = 40) then underwent 3 experimental sessions: fasting, baseline and isolation (the order of sessions was counterbalanced across participants) and subsequently an MRI scan with the cue-induced craving task. On the baseline day, participants also underwent a functional localizer task. Cue-induced craving task: Participants saw cues for social contact, food and control cues depicting flowers. After each block of cues (showing 3 images), participants rated their self-reported social craving (after social blocks), food craving (after food blocks) and how much they liked the flower pictures (after control blocks). Functional localizer: Participants memorized a set of 5 images prior to the scan (4 different sets of images were counterbalanced across participants). Immediately before the localizer task, participants were shown the memorized pictures again. During the task, participants saw either one of the memorized pictures or a novel picture indicating whether they would be able to win money or not.

All participants were within a healthy weight range (BMI mean (standard deviation(SD))= 22.8(2.2)), reported frequent social interactions (monthly interactions, mean=49.1(31.7); minimum=10) and close relationships (number of close relationships, mean=12.3(5.1); minimum=3). Participants reported relatively^21^ low levels of pre-existing loneliness (UCLA loneliness-scale, mean=33.2(6.3), maximum=47 out of 80).

## Subjective social craving can be evoked by acute objective social isolation

Could we experimentally induce the subjective experience of social isolation in human participants? Human loneliness is not a simple product of objective isolation: people can be alone without feeling lonely, or feel lonely even in a crowd^22^. Moreover, experimentally induced isolation would be brief, relative to the human lifespan, and for ethical reasons, human participants (unlike rodents) would be able to predict when the isolation would end. In all, the first challenge of this research was to develop an experimental induction of objective isolation that created the subjective experience of unmet social needs in human participants. To address this challenge, we had socially-connected healthy human adults spend ten hours (9 am to 7 pm) alone, with no social interaction and no other social stimulation (e.g. social media, email, fiction). We used self-report questionnaires to assess people’s resulting subjective experience of loneliness and social craving.

After ten hours of social isolation, participants reported substantially increased social craving (t(39)=5.00, p<0.001), loneliness (t(39)=5.17, p<0.001), discomfort (t(39)=5.57, p<0.001) and dislike of isolation t(39)=4.13, p<0.001), and decreased happiness (t(39)=-4.21, p<0.001) compared to when they started isolation. Thirty-six out of forty individual participants reported feeling more lonely after isolation. As expected, after ten hours of food fasting participants reported increased food craving (t(36)=17.40, p<0.001), hunger (t(36)=23.90, p<0.001), discomfort (t(36)=13.56, p<0.001) and dislike of fasting (t(36)=6.28, p<0.001), and decreased happiness (t(36)=-3.05, p=0.004) compared to when they started fasting. Thus, both forms of abstinence evoked craving for the specifically deprived need, along with general discomfort and decreased happiness. However, we note that social craving after isolation was more variable across participants than food craving after fasting (mean(SD) of craving ratings, after 10 hours: food craving=80.25(19.39); social craving=66.38(24.52), Figure 2; Levene’s test indicated unequal variances (F(76)=15.86, p<0.001).

**Figure 2.**
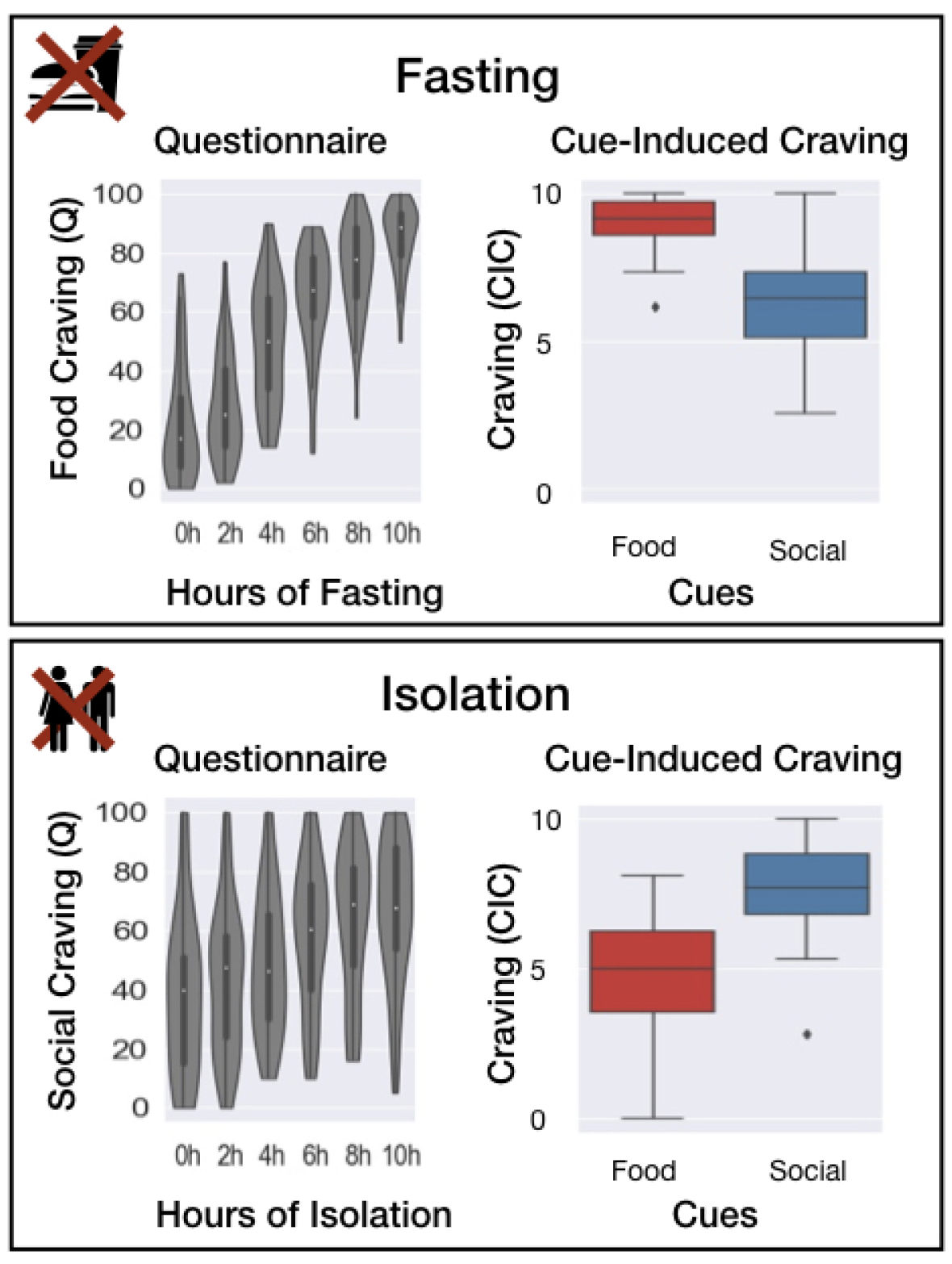
Behavioral results. The upper panel shows changes in self-reported food craving over time during fasting (left; n = 37) and in comparison to ratings of social craving during the cue-induced craving task (right; n =40). The lower panel shows changes in self-reported social craving over time during isolation (left; n = 40) and in comparison to ratings of food craving during the cue-induced craving task (right; n = 40). The boxplots in both panels indicate the median (dark center line), the interquartile range (IQR; box) and the 1.5 IQR minima and maxima (whiskers). Datapoints outside the whiskers are shown as individual data points.

## Following isolation, social cues evoke neural signatures of craving

In primates, aversive motivation (i.e., a negative state such as hunger or pain that motivates behavior to relieve the state^23^) is represented in the substantia nigra pars compacta and ventral tegmental area (SN/VTA^24^), and SN/VTA is activated by craving for food and for drugs of addiction^25–27^. In the substantia nigra in particular, ~70% of the neurons are dopaminergic, so fMRI signals in this brain area likely mainly reflect dopaminergic (DA) neuron activity^28^. We therefore hypothesized that acute isolation in humans might produce a social craving response in SN/VTA. Neuroimaging of the SN/VTA poses a technical challenge, though, because it is a small structure, adjacent to the sphenoid sinus (a large, air-filled cavity located anterior to the brainstem) and is therefore prone to distortions and signal loss^28^. To address this challenge, we optimized MRI image acquisition parameters, and used a newly developed atlas (see Methods section for details) to identify SN/VTA in individual participants’ brains. We also included an independent functional localizer task to identify voxels in each participant’s midbrain that are maximally sensitive to expected reward and novelty, consistent with DA activity^29^.

To measure dopaminergic midbrain responses to food cues and social cues, we developed a cue-induced craving (CIC) paradigm, in which participants viewed pictures of their favorite social activities, favorite foods, and a pleasant control (flowers). We included favorite foods so that we could compare social craving, within participants, to a well-established neural craving response, evoked by viewing food cues after several hours of fasting^26, for review^. In the anatomically defined SN/VTA (Figure 3a), responses to food cues were higher after fasting than after isolation (b=0.06, t=3.1, 95%CI=[0.0.02,0.09], p=0.002), but responses to social cues were not significantly higher after isolation than after fasting (b=0.006, t=0.7, 95%CI=[−0.01,0.03], p=0.50; for full results of the model including all main effects, see Table S1 in the Supplementary Materials (SM)). In the midbrain functional ROI (voxels maximally sensitive to reward and novelty, Figure 3b), responses to food cues were higher after fasting than after isolation (b=0.03, t=3.3, 95%CI=[0.01,0.05], p=.001), and responses to social cues were higher after isolation than after fasting (b=0.03, t=2.5, 95%CI=[0.006,0.05], p=0.01; see Table S2 in the SM for full results).

**Figure 3.**
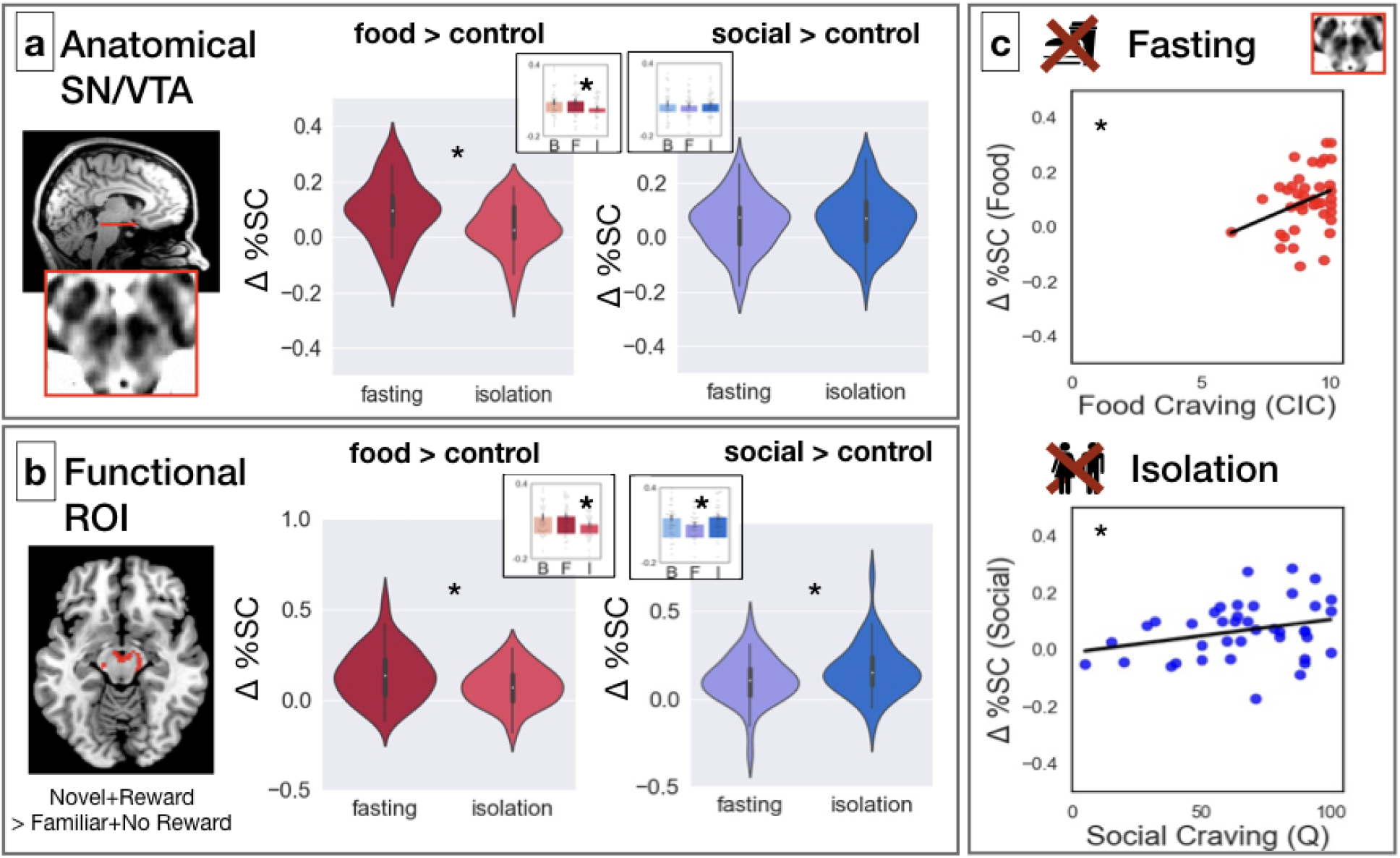
Univariate activity in response to food fasting and social isolation. Data (n = 40) shown for the anatomical ROI (a), the functional ROI (b) and correlations with self-reported craving (c). The violin plots depict the difference (in percent signal change) in response to food cues (contrast: food>flowers) and social cues (contrast: social>flowers) after fasting and isolation. The violins illustrate the distribution of the data, the white dots indicate the median, the bold dark grey vertical line the interquartile range (IQR) and the thin grey lines the 1.5 IQR minima and maxima. The insert barplots depict the contrast values for all three session: baseline (B), fasting (F) and isolation (I), showing the mean for each session. The error bars indicate the standard error of the mean. a: Responses in the anatomical SN/VTA were higher for food after fasting compared to isolation. b: Responses in the midbrain functional ROI were higher for food after fasting (compared to isolation) and for social cues after isolatio (compared to fasting). c: In the anatomical SN/VTA, responses to food cues were correlated with craving reported in the CIC task, following fasting (upper panel) and responses to social cues were correlated with craving reported on the final Questionnaire, following isolation (lower panel).

We also compared SN/VTA responses to food and social cues following fasting and isolation to responses to the same cue on the baseline day (when participants’ food intake and social interactions were not controlled or measured). Unexpectedly, when compared to the baseline day, deprivation led to a decreased response to the non-deprived cue, rather than an enhanced response to the deprived cue (see Figures 3 and S2). In the anatomically defined SN/VTA, responses to food cues were numerically but not significantly higher after fasting than baseline but significantly lower after isolation than baseline (b=-0.05, t=-2.1, 95%CI [−0.11,−0.003], p=0.038); responses to social cues did not significantly differ from responses at baseline for any of the sessions. In the midbrain functional ROI, responses to food cues were numerically but not significantly higher after fasting than and significantly lower after isolation than baseline (b=-0.06, t=-2.4, 95%CI [−0.12,−0.01], p=0.02); while social cues were numerically but not significantly higher after isolation than baseline and showed a trend to be lower after fasting (b=0.01, t=-1.8, 95%CI [−0.11,−0.004], p=0.07; see Tables S3 and S4 in the SM for full results) SN/VTA activity was correlated with self-reported craving, for both food and social cues. However, this correlation was significant for different measures in the two conditions. We measured self-reported craving in two ways: craving ratings during the task directly in response to the craving cues (*Food_Craving_CIC* / *Social_Craving_CIC)* and craving ratings on the questionnaire before participants went into the scanner (*Food_Craving_Q* / *Social_Craving_Q*). The two craving measures (*Craving_CIC* and *Craving_Q*) were correlated across participants in both conditions (food r(36)=0.52; p<0.001; social: r(38)=0.30; p=0.030); see Methods for details). We consider the two self-report measures to be estimates of the same hypothesized psychological process, and report results as significant after Bonferroni correction for two tests. After fasting, in the anatomical SN/VTA, the response to food cues (versus flowers) was positively correlated with the participant’s self-reported food craving measured during the cue-induced craving task (*Food_Craving_CIC*: r(38)=0.31; p=0.025; Figure 3c), but not with food craving on the final questionnaire (*Food_Craving_Q,* r(36)=0.05; p=0.375). Because the data for food craving measured during the cue-induced craving task is truncated at the upper limit of 10, we also analyzed the correlation using a truncated regression model (see Methods for details): SN/VTA activity in response to food cues remained associated with higher cravings ratings during the task (*Food_Craving_CIC*: b=0.04, t=2.10, p=0.038). After isolation, the SN/VTA response to social cues (versus flowers) was positively correlated with self-reported social craving on the final questionnaire (*Social_Craving_Q*, r(38)=0.36; p=0.011; Figure 3c); but not with social craving measured during the cue-induced craving task (*Social_Craving_CIC*: r(38)=0.054; p=0.370).

As a direct test of the similarity of activity patterns for food cues after fasting, and social cues after isolation, we implemented a multivoxel pattern analysis (MVPA). We trained a classifier on the pattern of activity in the SN/VTA after fasting to food cues and flower images. This trained classifier then generalized to above chance decoding (α=0.001; see Methods section for details and Figure 4 for an illustration of our multivariate analysis approach) of social cues from flower images after isolation (mean accuracy=0.542, bootstrapped-CI=[0.505,0.581], p<0.001) but did not show significant above-chance accuracy at baseline (mean accuracy=0.534, bootstrapped-CI=[0.497,0.573], p=0.007)). However, it is important to note that the classification accuracy for social cues did not differ between the isolation and baseline sessions.

**Figure 4.**
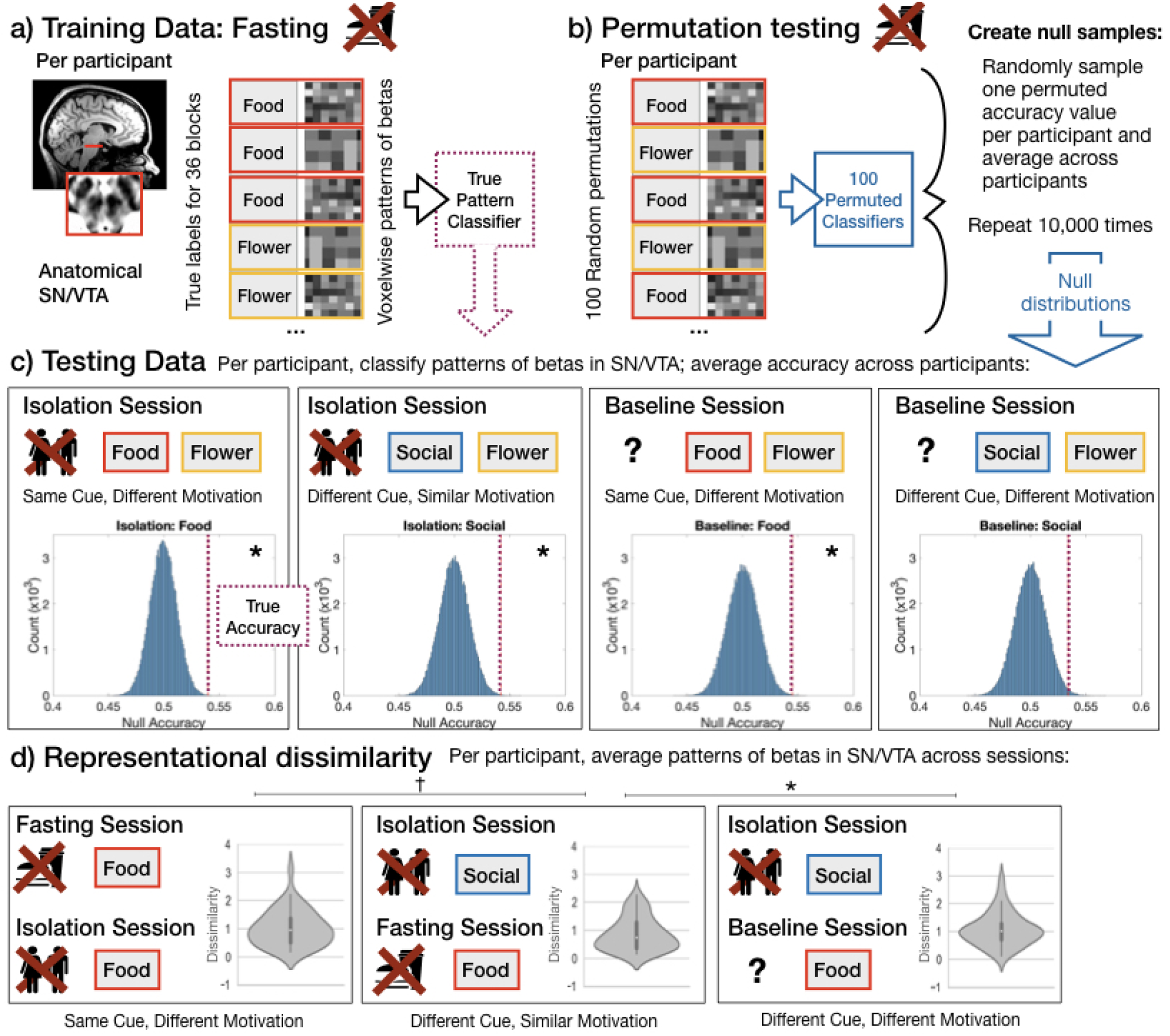
Multivoxel pattern analysis. (a) A linear classifier was trained to distinguish the pattern of activity across voxels in the anatomical SN/VTA of each participant (n = 40), in response to food and control cues after fasting. This classifier was then tested on patterns of activity in the same participant in the other two sessions. (b) For comparison, we generated null distributions using permutation and bootstrapping procedures: first, we permuted the labels within runs for each participant’s training data, and then tested the permuted classifier on each testing dataset. We created a null distribution by randomly sampling one of the permuted accuracies for each participant and then averaging the accuracy values across subjects. This procedure was repeated 10^5^ times to generate the group level null distributions. (c) Null distributions for each testing data set. The dotted red line is showing the true accuracy value (i.e., classification accuracy from the real classifier) for each testing dataset. The classifier generalized to (i.e. successfully decoded above chance) food cues (vs control cues) on both isolation and baseline days; and to social cues (vs control cues) on the isolation day, but not on the baseline day. (D) We directly compared the spatial pattern of activity for deprived and non-deprived cues. Note that these results show fisher-transformed dissimilarity, a measure of representational distance, so lower numbers indicate a more similar spatial pattern. The violins illustrate the distribution of the data, the white dots indicate the median, the bold dark grey vertical line the interquartile range (IQR) and the thin grey lines the 1.5 IQR minima and maxima. Social cues after isolation were more similar to food cues after fasting than to food cues at baseline. Food cues after fasting were (trending) more similar to social cues after isolation than food cues after isolation. Both (C) and (D) show that the pattern of activity in SN/VTA is determined not only by the category of visual stimulus, but by the motivational salience of the category, with craved cues evoking a similar pattern whether what is craved is food or social interaction.

In addition, the classifier was able to decode food from flower images after isolation and baseline (isolation: mean accuracy=0.540, bootstrapped-CI=[0.494,0.584], p<0.001; baseline: mean accuracy=0.545, bootstrapped-CI=[0.509,0.594], p<0.001). Given that the classification algorithm was always used to distinguish between a target stimulus and flowers, one might worry that the generalization is simply due to learning about flowers. We addressed this concern using a representational similarity analysis that directly compared food and social cues in deprived or undeprived states.

The pattern of SN/VTA activity in response to social cues on the isolation day was more similar to the pattern of food cues on the fasting day, than to food cues on the baseline day (mean fisher-transformed dissimilarity: social_craved-food_craved=0.87, social_craved-food_noncraved=1.09; t(39)=2.0, p=0.025). Indeed, SN/VTA responses to different stimuli in a similar motivational state (social_craved-food_craved) trended towards being more similar to each other than the responses to the same stimuli across different motivational states (food_craved–food_noncraved; mean correlation: 1.03; t(39)=1.4, p=0.08).

In sum, these results suggest that across all participants, SN/VTA shows an increased response to social cues after objective social isolation compared to after fasting, with a spatial pattern that is similar to the response to food cues when hungry. The magnitude of this response was variable across participants, and larger in those who reported more social craving after the acute isolation period. We predicted that individual variability in response to objective isolation might reflect pre-existing differences in participants’ social network size and/or chronic loneliness. Consistent with this prediction, participants with higher levels of chronic loneliness during the initial screening reported less craving for social contact in response to the social cues in the CIC task (*Social_Craving_CIC*: r(38)=-0.37; p=0.020) and somewhat less craving after 10 hours of isolation, on the online questionnaire (*Social_Craving_Q*: r(38)=-0.30; p=0.059). People with higher pre-existing chronic loneliness also showed a muted response in SN/VTA to social cues after acute isolation (r(38)=-0.33; p=0.036). Individual differences in social network size did not predict either self-reported or neural responses to acute social isolation (all p-values>=0.33). We explored whether pre-existing loneliness was associated with different responses to food cues after fasting and find that while loneliness did not affect self-reported food craving (p=0.431), higher loneliness was associated with a trend towards lower post-fasting SN/VTA responses to food cues (r(38)=-0.30; p=0.062).

### SN/VTA: Correlations with cravings

In our pre-registration, we planned to test whether the magnitude of SN/VTA response to a specific deprived cue (food or social cues) was correlated with self-reported craving for that cue. As reported above, these predicted correlations were observed. In addition to these pre-registered analyses, as prompted by a reviewer, we explored the broader structure of correlations between behavioral and neural measures of craving.

We tested whether the correlations between SN/VTA and self-reporting craving (on the cue-induced craving task, CIC) were restricted to the deprived day. We found that they were not. Self-reported craving for food was correlated with SN/VTA response to food cues on both of the non-fasted sessions (baseline session: r(38)=0.32; p=0.045; isolation session: r(38)=0.43, p = 0.006). Because the data for food craving measured during the cue-induced craving task is truncated at the upper limit of 10, we also analyzed the correlations using a truncated regression model (see Methods for details): For both non-fasted sessions, SN/VTA activity in response to food cues remained associated with higher cravings ratings during the task (*Baseline*: b=0.02, t=2.11, p=0.034; *Isolation*: b=0.02, t=3.00, p=0.003). Self-reported craving for social interaction was correlated with SN/VTA response to social cues on both of the non-isolated sessions (baseline session: r(38)=0.37; p=0.018; fasting session: r(38)=0.35, p = 0.027). Thus, individual differences in the magnitude of SN/VTA response to a cue are correlated with simultaneously measured self-reported craving for that cue, whether or not that cue has been selectively deprived.

## Mixed effects model: striatum

Although our primary hypotheses focused on the dopaminergic midbrain, and particularly SN/VTA, we also investigated responses in the striatum, a major target of projections from midbrain DA neurons^30^. A mixed effects regression model tested effects of cue (food, social, flowers) and session (fasting, isolation) for each subregion of the striatum separately (nucleus accumbens (NAcc), caudate nucleus (Ca), and putamen (Pu). We report results as significant at p<0.017 (0.05/3).

Food cues (compared to flowers) evoked a stronger response after fasting (compared to after isolation) in nucleus accumbens (NAcc, b=0.08, t=4.7, 95%CI [.05,.11], p<0.001; Figure 5) and in putamen (Pu, b=0.03, t=3.2, 95%CI [.01,.05], p=0.002) but not in caudate nucleus (Ca, b=0.025, t=1.9, 95%CI [−.001,.05], p=0.061). Social cues evoked a stronger response after isolation (compared to after fasting) in Ca (b=0.04, t=3.0, 95%CI [.01,.06], p=0.003) but not in NAcc (b=0.04, t=2.1, 95%CI [.002,.07], p=0.04) and Pu (b=0.02, t=1.4, 95%CI [−.007,.04], p=0.17). Similar to the SN/VTA, when compared to the baseline day, deprivation led to a decreased response to the non-deprived cue in striatal subregions, rather than an enhanced response to the deprived cue (an enhanced response to social cues following isolation compared to baseline in the caudate did not survive correction for multiple comparisons). For full results, see tables S5 – S12 in the SM.

**Figure 5.**
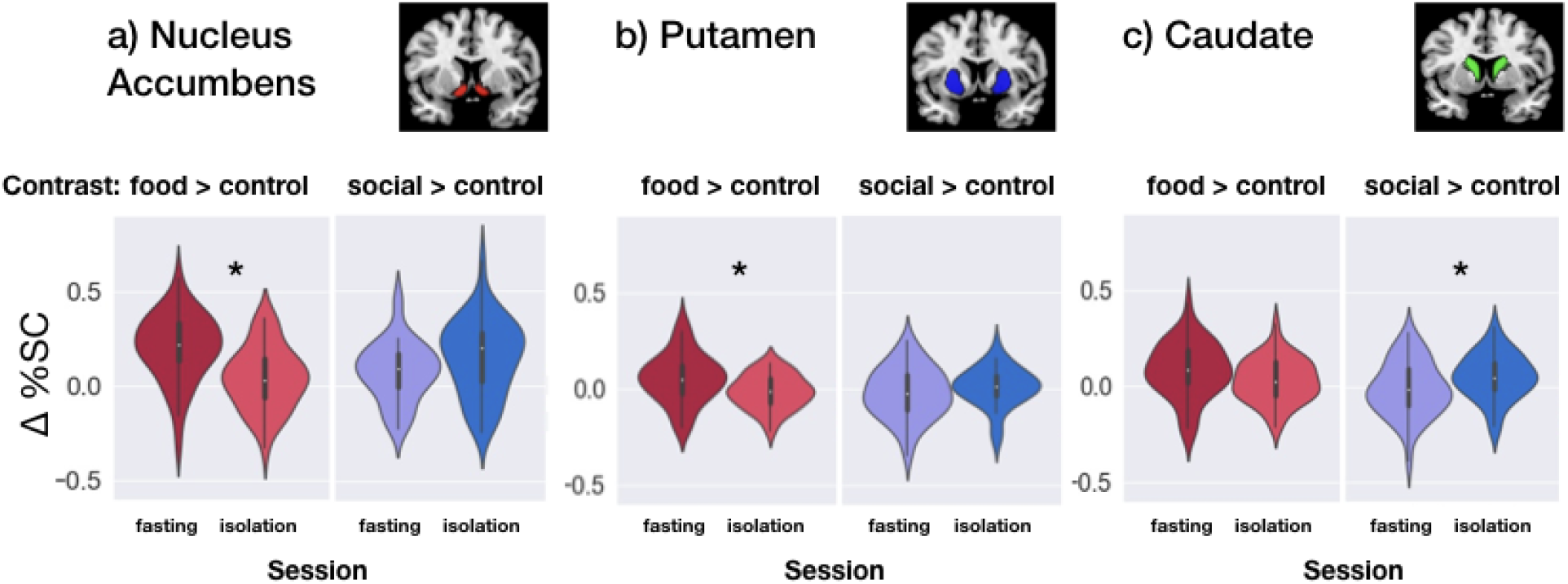
Univariate activity in response to food fasting and social isolation within the striatum. The striatum was divided into three subregions: Nucleus Accumnbens (a), Caucate Nucleus (b) and Putamen (c). The violin plots depict the difference (in percent signal change) in response to food cues (contrast: food>flowers) and social cues (contrast: social>flowers) after fasting and isolation. The violins illustrate the distribution of the data (n = 40), the white dots indicate the median, the bold dark grey vertical line the interquartile range (IQR) and the thin grey lines the 1.5 IQR minima and maxima.

Thus, while the responses in SN/VTA were similar and overlapping for food craving and social craving, responses in the striatum were dissociable. We conducted exploratory analyses to test whether subjective craving of food and social contact (*Craving_CIC*) was correlated with striatal activity for any condition or any of the three subregions (reporting results significant at α<0.017 to correct for multiple comparisons) and found no significant correlation in the striatum ROIs (all p-values>0.038).

## Exploratory analyses: Downstream targets of SN/VTA

We also conducted exploratory analyses testing the effects of food and social craving in other brain regions associated with craving. We selected four ROIs based on meta-analyses of craving across different modalities (see Methods section for details): amygdala, insular cortex, anterior cingulate cortex (ACC) and orbitofrontal cortex (OFC). A mixed effects regression model tested effects of cue (food, social, flowers) and session (fasting, isolation) for each region and we report results as significant at p<0.0125 (0.05/4), to account for the four regions tested.

Food cues (compared to flowers) evoked a stronger response after fasting (compared to after isolation) in ACC (b=0.07, t=2.9, 95%CI [.02,.12], p=0.005) and trending stronger responses after fasting in Insula (b=0.03, t=2.0, 95%CI [.0003,.07], p=0.052) and Amygdala(b=0.04, t=1.9, 95%CI [.0007,.07], p=0.055) but not in OFC (b=0.03, t=1.0, 95%CI [−.1,.04], p=0.337).

Social cues evoked a stronger response after isolation (compared to after fasting) in OFC (b=0.11, t=2.53, 95%CI [.03,.20], p=0.012) but in none of the other ROIs (ACC: b=-0.007, t=-0.3, 95%CI [−.06,.05], p=0.787; Insula: b=0.02, t=1.0, 95%CI [−.06,.02], p=0.324; Amygdala: b=0.02, t=1.0, 95%CI [−.02,.06], p=0.334). When compared to the baseline day, we find no significant effects in the exploratory ROIs. For full results, see tables S13 – S20 in the SM.

An exploratory whole brain random effects analysis yielded converging results: selective responses to food craving in nucleus accumbens, anterior cingulate cortex, periaqueductal gray, and amygdala, and selective responses to social craving in caudate nucleus, orbitofrontal cortex, and dorsomedial prefrontal cortex (see Tables S21 and S22 and Figure S4 in the SM). A conjunction analysis assessing overlapping activation between the food contrast (fasting > isolation) and the social contrast (isolation > fasting) did not show any suprathreshold voxels. We also calculated exploratory whole brain contrasts against baseline (i.e., food: fasting > baseline and social: isolation > baseline) but find no suprathreshold activity in this analysis. This result is in line with our ROI analysis in the SN/VTA showing similar magnitude of activity in in response to food cues following fasting and baseline and in response to social cues following isolation and baseline.

## Exploratory behavioral analyses

### Correlations between craving ratings

We explored whether individual differences in self-reported cravings for different cues are correlated with each other. We computed the Pearson’s correlation between each pair of average craving ratings (i.e., for food and social cues, measured after fasting, isolation and baseline). These correlations revealed some interesting structure (Figure S5). First, self-reported cravings for food and social interaction were moderately correlated at baseline (r=0.48), but less correlated after either need was deprived (mean r=0.14, z=1.64, p=0.05). Second, individual differences in cravings for social interaction were somewhat, but not significantly, more stable across the three sessions (mean r =0.33) than craving for food (mean r=0.06, z=1.26, p=0.10). Finally, the strongest correlation of all was between self-reported craving for food after fasting and for social interaction after isolation (r=0.64). These results suggest that there is a reliable individual difference in craving responses following deprivation, shared across both fasting and isolation, consistent with our observation of a common neural response to both modes of deprivation.

### Differences in craving measures

To explore the differences in the results for the two craving measures (Craving_CIC and Craving_Q) in more detail, we included the following exploratory analyses: 1) We assessed the correlation between these measures and found that for both, food and social craving the CIC and Q measures were correlated within session, across individuals (food r(38)=0.52; p<0.001; social: r(38)=0.30; p<0.032). 2) We tested whether the correlations between craving measures and SN/VTA activity was different for these two measures. We implemented two regression models, one for the fasting session and one for the isolation session. The interaction between SN/VTA activity and craving measure (i.e., Craving_CIC and Craving_Q) and its main effects were entered as predictors. We did not find a significant interaction of craving measure * SN/VTA activity for social craving ratings (b=37.65, t=1.03, 95% CI=[−35.44, 110.74], p=0.31) or for food craving ratings (b=3.16, t=0.11, 95% CI=[−58.63, 52.31], p=0.91). This suggests that for social craving and food craving the correlations between the two ways of measuring craving and SN/VTA activity were not significantly different.

### Craving ratings baseline

Self-reported food craving on fasting days and social contact craving on isolation days were both higher than on the baseline day (*Food_Craving_CIC*): (t(39)=7.94, p<0.001, two-tailed; *Social_Craving_CIC* (t(39)=4.15, p<0.001, two-tailed). Self-reported food craving on the isolation day was lower than on the baseline day (t(39)=3.59, p=0.001, two-tailed); but self-reported social craving on the fasting day was not lower than on the baseline day (t(39)=0.64, p=0.526, two-tailed). Figure S2 shows the craving ratings for each session including the baseline day.

### Associations between isolation and chronic loneliness

Participants who reported higher levels of chronic loneliness showed a muted response in SN/VTA to both social cues (r(38)=-0.46, p=0.003) and food cues (r(38)=-0.37, p=0.018; two-tailed) during the baseline scan.

Similar to responses after isolation (see above), on the baseline day participants higher in chronic loneliness reported less social craving immediately before the scan (*Social_Craving_Q,* r(38)=-0.40, p=0.011; two-tailed), though chronic loneliness did not affect craving ratings during the task (*Social_Craving_*CIC, r(38)=-0.15, p=0.367; two-tailed).

## Discussion

In humans, acute mandatory social isolation evokes a neural “craving” response to social cues. Midbrain regions showed selective responses to food cues after fasting and to social cues after isolation. SN/VTA activity was higher in people who self-reported wanting food or social interaction more, following deprivation. The multivariate pattern of SN/VTA response was similar for food and social interaction when craved. People who are forced to be isolated crave social interactions similarly to the way a hungry person craves food.

Our findings are consistent with results from the mouse showing that dopaminergic neurons in the midbrain represent the neural substrate of social isolation^16^. In mice, dopaminergic neurons in the midbrain appear to encode an aversive “loneliness-like” state that motivates social engagement. Our findings suggest that there is a similar mechanism underlying social craving in humans.

Despite the fact that isolation lasted only ten hours, and the participants knew exactly when it would end, participants reported more loneliness and social craving at the end of the day than they did at the beginning. For people who are highly socially-connected, a day of social isolation is a large deviation from typical rates of social interaction. Although when chosen intentionally, solitude can be restful and rejuvenating^31,32^, the externally mandated isolation was subjectively aversive.

Our primary, pre-registered hypotheses concerned activity in SN/VTA. SN/VTA includes almost exclusively dopaminergic neurons, which respond with phasic firing to motivationally relevant cues (see^28^ for review) and is activated by craving for food and for drugs of addiction^25–27^. Using a cue-induced craving task, we found that SN/VTA responded more to food cues after fasting and more to social cues after isolation. That is, although participants reported general discomfort and reduced happiness after both fasting and isolation, SN/VTA responses were selective to the deprived cue. The magnitude of SN/VTA response varied across participants, and was correlated with self-reported craving for the corresponding cue: SN/VTA responses to food cues were correlated with self-reported craving for food (in all sessions), and SN/VTA responses to social cues were correlated with self-reported craving for social contact (also in all sessions). These results fit with the intuitive prediction that the deprivation of a need causes increased craving for the specific need^33^. The specific cravings evoke a generalizable pattern of activity in SN/VTA: patterns of response in SN/VTA in response to food when hungry were more similar to the pattern of response to social cues when isolated than to responses to food when sated. Thus, a common signal at the core of the “craving circuit” in the SN/VTA responds selectively to motivationally salient deprived cues, independent of their specific content.

By contrast, food and social craving led to dissociable responses almost everywhere else in the brain. Both whole brain and exploratory ROI analyses revealed that food craving evokes selective responses in anterior cingulate cortex and (less reliably) in insula and amygdala but not in orbitofrontal cortex, while social craving evokes selective responses in orbitofrontal cortex but not anterior cingulate cortex, insula or amygdala. Even in the striatum, the major target of dopaminergic projections from SN/VTA, craving for food and social contact were spatially dissociated. Fasting enhanced responses to food cues mostly in nucleus accumbens, whereas isolation enhanced response to social cues mostly in the caudate nucleus. Caudate nucleus activity during social craving is consistent with prior evidence of caudate activity when people re-live experiences of rejection by a significant other^34, for meta-analysis^. Though social rejection (being deliberately and specifically excluded from social interaction) is conceptually distinct from social isolation (being unable to access social interaction), both rejection and isolation could lead to increased social motivation^8^.

Food craving and social craving thus evoke both shared (SN/VTA) and unshared (striatum, cortical regions) neural responses. One open question concerns how these two kinds of craving might interact. That is, how does inducing food craving affect social motivation, or vice versa?

On one hand, there is some evidence that deprivation of one need leads to narrowed focus on the deprived need to the exclusion of other needs. An unexpected finding of our study was that, compared to the baseline (no deprivation) session, neural activity for the non-deprived need was *decreased*, rather than activity for the deprived need being selectively *increased*. This pattern of narrowing activity was observed for both food and social cues. For example, the midbrain response to food cues was highest after fasting, slightly but not significantly lower at baseline, and lowest after social isolation and eating to satiety. Putamen and nucleus accumbens responses to food cues were the same: not significantly higher after fasting than baseline, but significantly lower after isolation. These results fit with previous fMRI studies that find increased activity in midbrain and the “craving circuit” (i.e., striatum, OFC, ACC, amygdala and insula) in response to food cues after fasting compared to carefully induced satiety (e.g.,^25,35,36^), but not reliably when compared to baseline (e.g.,^37–39^). Here we observed a similar pattern for social craving: midbrain responses to social cues were highest after isolation, slightly but not significantly lower at baseline, and lowest after fasting. Thus, at baseline participants’ motivation may be spread across multiple sources of reward, and specific acute deprivation may serve to narrow and focus the brain’s motivational responses to the deprived target, rather than to enhance it. Depriving one need might thus reduce motivation to pursue other needs. Indeed, there is some evidence that people are less prosocial when hungry^40^, consistent with a reduction in social motivation caused by acute hunger, although see^41^.

On the other hand, there is also some evidence that deprivation of one need leads to increased motivation to pursue other sources of reward. In animal models social isolation can cause increased food consumption^42, for review^, increased susceptibility to substance addictions, and other general changes in motivational systems^8, for review^. That is, deprivation of social needs can result in generalized reward-seeking behavior, potentially as a form of compensation.

These two possibilities are not mutually exclusive: deprivation could lead to narrowed focus (reduced pursuit of other needs) or compensation (increased pursuit of other needs), depending on the duration and developmental timing of the deprivation. For example, short term acute isolation in adults may cause a temporary narrow focus on social connection, whereas long-term or developmental isolation could result in a shut-down of these adaptive efforts resulting in social withdrawal and other compensatory changes in non-social motivation^43^. In our study, people who reported higher levels of pre-existing chronic loneliness showed reduced activity in the SN/VTA in response to social cues, consistent with the idea that chronic isolation can lead to social withdrawal (^18,44^ but see^19,20^). However, the causal mechanism underlying the correlation with chronic loneliness remains unclear: it might as well be that individuals with generally reduced sensitivity of motivational brain areas are more prone to becoming lonely. Indeed, people reporting high chronic loneliness in our study also had lower SN/VTA responses to food cues. Chronic loneliness could thus be a consequence, rather than a cause, of general low responsiveness in SN/VTA.

In addition, the effects of chronic loneliness on social approach motivation might also be mediated via other factors affecting approach behavior more broadly, such as anxiety or depression.

This ambiguity of correlational observations highlights the importance of our design, experimentally inducing acute isolation to disentangle direct effects of isolation per se from individual differences in reward seeking behavior more generally.

There are two key limitations in this study that could be addressed in future research. First, we measured craving using passive viewing of craving-related cues and self-report measures of craving, rather than a direct test of motivation, such as participants’ willingness to expend effort or money to fulfill a need. Responses in the passive cue-induced craving task could also be influenced by low-level processes like increased visual attention to the deprived category (though the robust prior literature in both humans and animals makes visual attention an unlikely explanation of SN/VTA engagement^28, for review^). We used two different measures of self-reported craving, and while these were correlated with each other, different craving measures were significantly correlated with SN/VTA activity for food craving and to social craving. Thus, future studies should investigate the changes in subsequent behaviours that are predicted by SN/VTA responses following acute social isolation. Note though that also in domains like drug addiction and hunger, researchers have struggled to establish a gold-standard behavioral metric of subjective craving with better external validity than self-report^45^.

A second limitation is that we studied a small sample (n=40) of healthy well-connected young adults, mostly students, in one cultural context. Chronic loneliness disproportionately affects people older (i.e., the elderly^46^) and younger (i.e., adolescents^14^) than the participants studied here; future studies should test whether the current results generalize to those more vulnerable populations. Yet despite our restricted sample, we observed substantial variability between individuals at baseline, in both self-reported social craving and in SN/VTA response to social cues. We did not measure people’s social behaviours at baseline (e.g., number and quality of social interactions) or other traits broadly relevant to social interaction (e.g., trait extroversion, depression, anxiety). Future studies should investigate how social craving and SN/VTA responses to social cues respond to changes in people’s social environment over both short (hours) and medium (weeks or months) time scales, within individuals^47^.

In all, our finding of more selective SN/VTA response to social cues after isolation, as well as to food cues after fasting, fits the intuitive idea that positive social interactions are a basic human need, and acute loneliness is an aversive state that motivates people to repair what is lacking, similar to hunger. Thus, our research provides empirical support in human participants for the “social homeostasis” hypothesis developed based on animal models^43^. Despite differences in the duration and setting of social isolation, and in the anatomy of DA midbrain structures, both humans and mice seem to show midbrain craving responses for social interaction, as well as for food. Even this broad similarity of neural responses in mice and humans is encouraging for the translational prospects of mouse models of mental health disorders that affect social motivation – for example autism spectrum disorder^48^.

A vital question is how much, and what kinds of positive social interaction are sufficient to fulfill our social needs and thus eliminate the neural craving response. Technological advances offer incessant opportunities to be virtually connected with others, despite physical separations. Yet, some have argued that using social media only exacerbates subjective feelings of isolation^49, for review^ but see^50^. The potential for virtual interactions to fulfill social needs is particularly relevant when large populations are required to self-isolate, for example during a global pandemic. In early 2020, millions of humans experienced a sudden externally mandated period of relative or complete physical isolation from others, as public health officials sought to slow the spread of an infectious novel corona virus. This unprecedented upheaval in people’s social routines emphasized the need for a better understanding of human social needs and the neural mechanisms underlying social motivation.

## Methods

A Life Sciences Reporting Summary can be found in the supplementary materials.

### Participants

Participants (n=40) were healthy right-handed adults, ranging in age from 18-40 years (mean age 26 years; N=27 female). An a priori power analysis in G*Power 3.0^51^ targeted on the detection of medium effects (d=0.5, α=0.05 and 1-β=0.80) suggested a sample size of n=34. The targeted effect size was chosen based on findings from studies employing cue-induced craving paradigms for drug craving^52^, food cravings^25,37,53,54^ and internet gaming craving^27^ which report medium to large effect sizes in cue reactivity^52^. We performed power calculations for medium effects because social craving might be less intense or more variable than cue reactivity in drug craving and food craving. We therefore recruited 42 participants to account for potential attrition or exclusion for MRI data quality; two participants were unable to complete all experimental sessions and so were dropped from analysis, leaving 40 complete datasets.

Participants were recruited via e-mail lists and through online advertisements and flyers. Interested individuals filled out a screening questionnaire to assess eligibility for the study (questionnaire data were collected using REDCap Software, Version 5.5.17). People were eligible if they reported a healthy Body Mass Index (BMI: 16-30), no current calorie restricting diet, no permanently implanted metal in their body, no history of brain damage, and no currently diagnosed mental health disorder or substance abuse. Because we aimed to study social motivation in a sample of adults who have frequent and regular social interactions, we also excluded people who i) lived alone, ii) reported current feelings of loneliness on the UCLA Loneliness Scale^21^ (i.e., we excluded people with scores above 50, which is one standard deviation above the mean for a student sample^21^); or iii) reported smaller social network sizes than typically expected of adults^55^ according to a social network size measure^56^ and the Social Support Questionnaire^57^ (i.e., we excluded people with social networks 2 or more SD below mean, based on prior measured distributions from Von der Heide et al. 2014^58^). All experimental procedures were approved by MIT’s institutional review board, COUHES (couhes.mit.edu). Participants signed a consent form describing all experimental procedures before participating in the study. Each participant was compensated with $350 for participating in three fMRI sessions and completing the 10 hours of food fasting and 10 hours of social isolation.

### Experimental Procedures

Each participant was scanned in three fMRI sessions, separated by at least 24 hours. Figure 1 shows an overview of the experimental procedures. One session followed 10 hours of food fasting; one session followed 10 hours of social isolation, and one session was a baseline without any mandated prior abstinence. All participants underwent all the three experimental conditions (baseline, food fasting and social isolation). Between participants, the order of sessions was counterbalanced; each participant was pseudo-randomly assigned to one of the possible orders of the three different sessions with the restriction that all 6 possible sequences were approximately equally likely in the full sample. Data collection was not performed blind to the conditions of the experiments.

#### Food fasting

Participants were asked to abstain from consuming any food or drinks/coffee (except water) for 10 hours before the fMRI session. We scheduled each fMRI session at 7pm in the evening; thus, participants were asked to refrain from eating after 9am on the day of the fasting session. We followed methods of previous food craving studies (for review:^26^) in which participants are instructed to fast at home. Fasting was confirmed through self-report upon arrival. We also asked participants to abstain from all forms of exercising on the day of food fasting in order to avoid exhaustion. Participants filled out an online questionnaire, rating their momentary food craving, hunger, discomfort, happiness and dislike of fasting (on a visual analog scale (VAS) anchored at 0 (not at all) to 100 (extremely)), every two hours during the food fasting period. Before the scan, participants were asked to select a meal that they would receive following the scan on an online food ordering platform (Grubhub). Participants selected their food while they were still fasted, and we gave no restrictions for what or how much to order.

#### Social isolation

Participants were socially isolated for 10 hours. On the day of the isolation session, participants arrived at the McGovern Institute for Brain Research, MIT building 46, at 8.15am. Because we aimed to keep all social interactions between the social isolation and the fMRI scan to a minimum, participants were given extensive instructions about the paradigm and MRI session, and a mock scanner session, before starting social isolation. Subsequently, participants gave their phones and laptops to the experimenter and were guided to a room containing an armchair, a desk and office chair, and a fridge with a selection of food, snacks and beverages. Participants remained in that room from 9am until 7pm. In advance of the session, participants were invited to send us text documents (without any social content) to read or work on during isolation; approved documents were printed or transferred to the provided laptop. In addition, we provided puzzles, Sudoku, coloring pages, non-social games (e.g., Tetris, Bubble Shooter, etc.) and drawing/writing supplies. Participants were provided with a laptop (with parental controls enabled), allowing them to visit only our Slack channel (i.e., an online messenger software allowing communication between a group of people [www.slack.com]) and the webpage containing our online questionnaire. Messaging in Slack was restricted to informing participants about the arrival of food delivery and for emergencies (i.e., in case participants ran into problems which required assistance from the research team during isolation). Participants filled out an online questionnaire rating their momentary social craving, loneliness, discomfort, happiness and how much they disliked isolation (VAS anchored at 0 (not at all) to 100 (extremely)) every two hours during the social isolation period. The fMRI session was conducted immediately after the social isolation. Following the scan, a member of the research team chatted with the participants about their experiences during isolation and made sure participants were not feeling troubled. Because living in a shared household was a prerequisite for participating in the experiment, participants were expecting to meet with at least one other person after the experiment.

#### Baseline

Participants came into the lab at 7pm and completed the same fMRI tasks as in the other two conditions (in addition to a functional localizer task, see below). Participants were asked to not be hungry at the time of the scan.

### fMRI

Participants were in the scanner for around 1 hour in each session. We started with anatomical scanning. For each participant, structural whole-head T2*-weighted structural images were collected in 176 interleaved sagittal slices with 1 mm isotropic voxels (FOV: 256 mm). In addition, whole-head T1-weighted structural images in 176 interleaved sagittal slices with 1 mm isotropic voxels (FOV: 256 mm) were collected. The T2* weighted anatomical scan was collected for anatomical identification of midbrain nuclei (i.e., the high content of iron in SN/VTA and red nucleus makes the T2* shorter and darker in these areas ^28,59^). We confirmed the identification of mid-brain structures by registering to the newly available atlas of subcortical nuclei from Pauli et al. ^60^ and defined separate regions of interest in the dorsal and ventral striatum. We also collected a field map (phase-difference B0 estimation; TE1=3.47ms, TE2=5.93ms) to control for spatial distortions, which are particularly problematic in midbrain fMRI ^28,59^. During acquisition of the anatomical images and the field map (~15min in total) participants lay quietly in the dark.

Subsequently, we collected functional data during six runs of a cue-induced craving task (see below for details). Each functional run consisted of 147 volumes with 58 T2*- weighted echo planar slices (EPIs; TR=2000 ms, TE=30 ms, FoV=210 mm, 70×70 matrix, yielding a voxel size of 3×3×3 mm) acquired as a partial-head volume in an A-P phase encoding direction using interleaved slices. The scanning parameters were extensively piloted (N=11) using the functional localizer task (see below) and the parameters showing the best signal-to-noise ratio (SNR) were selected for the study. Despite the small structure of the SN/VTA, we thus chose 3mm isotropic voxels because of their higher SNR compared to smaller voxels^61^. The angle of the slices was approximately 20 degrees away from the plane of the standard anterior commissure-posterior commissure (AC-PC) to avoid placing SN/VTA and the sphenoid sinus in the same slice plane. This reduced geometric distortion to the point that standard distortion correction methods could be applied^28^. The cue induced craving paradigm took approximately 26 minutes total.

#### Cue induced craving (CIC) task

We designed a novel CIC task to simultaneously measure craving for food and for social interaction, relative to a control. Participants viewed colored images depicting: i) groups of individuals as they meet, talk, laugh, smile, etc.; ii) different kinds of highly palatable foods such as cake, pizza, chocolate, etc., iii) attractive flowers as the control condition.

On each trial, participants saw a single photograph and 3-5 word verbal description, for 5 sec. The combination of visual and verbal cues was intended to maximize deep semantic processing of the relevant attributes. Each trial was followed by a 1s rest period (during which a fixation cross was displayed). Three consecutive trials were presented in a block of the same condition (food, social, control). Each block was followed by a jittered 2-6 second rest period. Subsequently, participants self-reported how much they were currently craving food (on food blocks) or social interaction (on social blocks). After control blocks, participants rated how much they liked the flower image, in order to match the demand for response preparation. A second jittered 2-6 rest period preceded the onset of the next block. In total, participants saw 18 blocks (54 trials) per condition, per scan session. The trials on each day were unique, so in total participants saw 36 blocks (108 unique images with descriptions) per condition. The duration of the task was approximately 30 minutes – divided into 6 runs, each run had a duration of approximately 5 minutes.

The stimuli for the CIC task were tailored to each individual’s preferred foods and modes of social interaction. During the initial screening, participants were asked to list their top ten favorite foods and social activities. Stock photographs illustrating these specific foods and activities were selected from a large public database (https://www.pexels.com/), and then verbal labels were added using the participant’s own descriptions. Food descriptions included “fluffy syrup-drenched pancakes”, “creamy cheesy macaroni”, “refreshing mixed fruit salad”, and “yummy vanilla cake with sprinkles”. Social descriptions included “chatting and laughing together”, “joking around with friends”, “supporting each other through workouts”, “enjoying a conversation together.” Social pictures were all matched for gender of participants (i.e., for a male participant, all social photographs included at least one man). The stimuli were images of strangers, rather than images of the participant’s own friends and family, in order to (i) match the food and control images for novelty, since SN/VTA activity is sensitive to novelty^62^, (ii) match image quality across conditions and participants, and (iii) avoid unmeasured variability in the quality or current status of participants’ relationships with specific individuals. Control trials presented attractive photographs of flowers accompanied by positive valence verbal descriptions (Figure 1). For group-level results in response to cues (across sessions), see Table S23 and Figure S6 in the SM.

#### Functional localizer task

During the baseline session, each participant completed a functional localizer at the end of their scan. We anticipated that anatomical localization of SN/VTA might be difficult, given the strong susceptibility to magnetic distortions in the midbrain^28,59^. We therefore designed a task to functionally identify relevant midbrain regions in each participant individually. The task was an adapted version of Krebs et al. 2011^62^. Because midbrain dopaminergic neurons in SN/VTA respond to both novelty and reward^62–64^, we contrasted novel and rewarding stimuli against familiar and non-rewarding stimuli. However, our pre-registered hypotheses focus on the anatomical localization strategy, so we treat analyses of the functionally localized regions as exploratory.

A potential limitation of this approach is that it was targeting voxels responding to secondary reinforcers (i.e., money) instead of primary reinforcers (such as food and social contact). In animal models, dopamine neurons in the midbrain exhibit burst firing in response to both, primary reinforcers and also to conditioned stimuli once the conditioning is established (e.g., ^65^ and for review:^66^). Conditioned stimuli are somewhat less effective than primary rewards in terms of response magnitude and fractions of neurons activated^67^. Thus, by localizing SN/VTA based on activity in response to anticipated financial rewards, we may have identified only a subset of voxels that respond to primary rewards. Importantly, both food and social cues would be similarly affected, so our key claim of similarity between these responses would not be undermined. In addition, our functional localization method was complementary to an anatomical ROI, which would not be affected by this issue.

Before beginning the localizer task, participants memorized a set of 5 images depicting abstract art (all images taken from the free stock pictures site (https://www.pexels.com/)). During the task, the abstract art images served as cues to the condition of the current trial. The task had two conditions: a reward/loss condition (*reward*) in which participants could earn or lose money depending on whether their responses were correct and fast enough, and a non-reward condition (*non-reward*) in which participants always received $0 regardless of their response. Each trial started with an abstract art image. The previously memorized (*familiar*) images indicated a *non-reward* trial. Abstract art images that were not previously observed (*novel*) indicated a *reward* trial. After the cue, participants saw a number between 1-9 (excluding 5) for 100ms on the screen. Their task was to press an assigned button indicating whether the number is below or above 5 as fast as possible. Initially, correct responses were required in less than 500ms; after 10 consecutive correct answers, this window was reduced to 400ms. After they pressed the button, participants saw the outcome indicating whether they won $1 (*reward* trial, correct response, fast enough), lost $0.20 (*reward* trial, wrong response or too slow), or received $0 (*non-reward* trial). In total, participants played 80 trials (40 trials per condition) and the duration of the task was approx. 10 minutes. Participants responded correctly and within the time limit on 87% of *reward* trials and 69% of *non-reward* trials. The earnings from this task were added to participants’ compensation after the baseline session. This design allowed us to compare responses to novel stimuli predicting reward versus familiar stimuli predicting no possibility of reward. For group-level results within the midbrain in response to reward > non-reward, see Figure S7 in the SM.

### Behavioral data analysis

#### Questionnaire data

For each participant we collected two measures of social network size (i.e., number of monthly interactions and number of close relationships). These scores indicate the size of participant’s social network on different hierarchical levels^55^. However, because the measures were highly correlated (r(39)=0.58, p<0.001), we z-transformed and averaged the two measures for each participant. This gave us an indicator of participant’s social network size relative to the sample’s average social network size. In addition, we calculated a loneliness score for each participant using the UCLA loneliness scale^21^. We tested how self-report ratings of hunger, food craving, discomfort, happiness and dislike of fasting provided during food fasting changed over the course of 10 hours using paired t-tests comparing the first rating (collected at beginning of fasting) and last rating (collected after 10 hours of fasting). We used the same analysis for the ratings provided during social isolation: loneliness, social craving, discomfort, happiness and dislike of isolation. Data distribution was assumed to be normal but this was not formally tested. Three participants missed filling out the the first or last round of the online questionnaire during fasting, so statistical analyses involving this questionnaire were conducted with a sample size of n = 37.

### fMRI Data analysis

#### Preprocessing

We used open source preprocessing pipelines for fMRI data, developed through the nipy and nipype^68^ initiatives. We used the heudiconv python application which uses dcm2niix to convert raw scanner data into the NIFTI image file format, then organizes that data into a BIDS-formatted directory structure. The FMRIPrep application^69^ was used to minimally preprocess the anatomical and functional data (using default settings but including susceptibility distortion correction using fieldmaps (see below)). Using FMRIPrep, we skull-stripped anatomical images first roughly using the atlas-based ANTS program^70^, and then refined it using information from Freesurfer surfaces after reconstruction was completed^71^. Brain tissue segmentation was performed with the FMRIB Software Library (FSL) FAST program^72^. Images were spatially normalized to 2mm isotropic MNI-space using the multiscale, mutual-information based, nonlinear registration scheme implemented in ANTS. We visually inspected brain masks, tissue segmentation and freesurfer surfaces. Susceptibility distortion correction was performed using phase-difference B0 estimation^73^.

A reference image for each run was generated from the input BOLD timeseries. A functional brain mask was created using a combination of FSL, ANTS, AFNI and nilearn tools^74^. Using FSL’s MCFLIRT program^75^, we estimated and corrected for head motion, resulting in a coregistered BOLD series as well as motion-based confound regressors. Any run containing a framewise displacement greater than 0.4 mm on more than 25% of the total frames was excluded from additional analyses. Additional confound regressors were generated, including other measures of motion (framewise displacement and DVARS and anatomical CompCor^76^ timeseries derived from CSF and white matter tissue segments). The reference image of each run was aligned with the anatomical image using FreeSurfer’s program “bbregister”^77^. The timepoint-to-functional reference transformation, the functional reference to anatomical transformation, and the anatomical-to-MNI transformation was concatenated into a single transformation and used to transform each functional timeseries into MNI template space. Spatial smoothing was performed on the FMRIPrep outputs with a 6mm smoothing kernel using FSL’s SUSAN tool^78^, which uses segmentation boundaries to avoid smoothing across tissue types. MRIQC, an opensource quality assurance software tool^79^, was used to generate additional reports which display Image Quality Metrics (IQMs).

#### Modeling

Analyses were conducted using the nipype framework^68^. For run-level analyses, the preprocessed timeseries was assessed with algorithms from the Artifact Removal Toolbox (ART)^80^ to identify frames within the run that have an abnormal amount of motion (0.4 mm of total displacement, or an intensity spike greater than 3 standard deviations from mean). The design matrix included boxcars for the experimental conditions convolved with a double-gamma hemodynamic response function (HRF), and nuisance regressors representing frame-wise motion, the anatomical CompCor regressors derived from white matter and CSF, as well as impulse regressors for volumes identified by ART. A high-pass filter (120 Hz) was applied to the design matrix and the smoothed data. The model was evaluated using FSL’s FILM program^81^. Subject-level contrast maps were generated using FSL’s FLAME^81^ in mixed-effects mode.

#### Data exclusion

Exclusion criteria were pre-established in our pre-registration (https://osf.io/cwg9e/). We excluded 3 runs of fMRI data from the overall sample (from 2 participants from the Cue Induced Craving task) based on the following criterion: Any run containing a framewise displacement greater than 0.4 mm on more than 25% of the total frames will be excluded from additional analyses.

#### ROI definition

We included functional voxels which overlapped at least 75% with the substantia nigra pars compacta (SN) and the ventral tegmental area (VTA) region (number of voxels=161) from the probabilistic atlas of human subcortical nuclei^60^. Because the striatum is a major target of projections from midbrain neurons and their firing results in increased DA transmission in the striatum^30,82,83^, we expected to see the same pattern of results in the striatum, i.e.: increased activation to food cues after food deprivation and to social cues after social isolation and a positive correlation between activity in striatum and self-reported craving (for both, food and social craving). Thus, we also included 3 additional ROIs in our analysis: putamen (Pu; number of voxels = 2530), Caudate Nucleus (Ca; number of voxels = 2523) and Nucleus Accumbens (NAcc; number of voxels = 300) also using the probabilistic subcortical atlas^60^.

#### Functional ROI definition

To define subject-specific ROIs, we used individual activations of each participant in the localizer task. The fMRI time series were analyzed using an event-related design approach implemented in the context of the GLM. The model contained two regressors separately modeling the presentation of novel/reward cues, and familiar/non-reward cues (i.e., when the abstract art images were presented, 2s). We also included one regressor for the time period of button press and outcome (1.1s). Because we did not add any jitter between button press and presentation of outcome (as this was not the contrast of interest), we modeled the whole segment as one block. For each participant, we calculated the target contrast *novel reward* > *familiar non-reward.* We then used a mask encompassing the whole midbrain as the search space for the selection of individual voxels. In each participant we selected the top 100 active voxels within the search space in response to the target contrast.

#### Spatial and functional differences in anatomical and functional ROIs

We explored the overlap in voxels between functional and anatomical SN/VTA ROI in each participant and found that it was variable across participants, with the maximum overlap being ~1/3 of the voxels. In the SM (Figure S8), we show a histogram of the overlap between the two masks for acros subjects. On the other hand, these two methods are likely measuring a similar underlying function because the mean activity in response to social cues following isolation was highly correlated across subjects (r(38) = 0.81; p < 0.001), as shown in the scatterplot in Figure S9 in the SM. Thus, we interpret the difference in results between functional and anatomical mask as reflecting the uncertainty of the measurement.

#### Definition of exploratory ROIs

We selected ROIs based on converging results from five meta-analyses of craving across different modalities^26,84–86^ identifying signals of craving in orbitofrontal cortex (OFC), amygdala, anterior cingulate cortex (ACC) and insula. Because the meta analyses report varying coordinates for the foci of activation, we chose to create ROIs based on an anatomical atlas instead (i.e., Harvard-Oxford cortical and subcortical probabilistic anatomical atlases; included in FSL). We selected all voxels that showed a minimum probability of 50% for being in the respective area and extracted the mean activity from those voxels in response to food, social and control cues for each session (fasting, baseline and isolation). We then calculated a mixed model regression for each region within this “craving circuit” (and report results as significant at p < 0.0125 (0.05/4)). In addition, due to previous findings showing that ventral striatum and nucleus accumbens are activated during craving for food and drugs^85^ and associated with altered responses in chronic loneliness^18,20^, we explored this region further at the suggestion of a reviewer by selecting the top 100 voxels within the NAcc that were active in response to the midbrain localizer task (i.e., active in response to reward anticipation) and assessed whether these voxels code for food and social craving (see SM for results).

#### Univariate Analyses

For our planned analyses, we used mixed effects regressions (using Matlab 2019b’s *fitlme* function) to estimate the fixed effects of cue, deprivation session, and the critical interaction of cue and deprivation session, on response magnitude in the ROIs, controlling for each session’s average framewise displacement (i.e. head motion), with subject included as a random effect. Following the recommendation of Barr, Levy, Scheepers, & Tilly, 2013^87^, we used maximal models, including participants as a random effect with both random intercepts and random slopes. Data distribution was assumed to be normal but this was not formally tested. We modeled these effects in the anatomically defined SN/VTA (pre-registered analysis) and in the functionally defined ROI of voxels maximally sensitive to reward and novelty (exploratory analysis). Thus, for each ROI, we tested two mixed effects models:

1. <u>Target comparison: Fasting vs. Isolation</u> First, we tested our key hypothesis, that responses to food cues (relative to flowers (control)) would be higher after fasting, and responses to social cues (relative to control) would be higher after isolation. In this model, the fixed effects were: session (fasting vs isolation; contrast coded), cue (control, +food, +social; indicator coded), the interactions of cue and session, and mean framewise displacement in that session; we also included random intercepts and random slopes for each participant.
2. <u>Exploratory analyses: baseline session</u> To test whether responses were different on the deprivation days, compared to the baseline day, we modeled responses on all three sessions by including fixed effects of session (baseline, +fasting, +isolation; indicator coded), cue (control, +food, +social; indicator coded), the interactions of cue and session, and mean framewise displacement in that session; we also included random intercepts and random slopes for each participant.

The command for both models was: *fitlme(Data,’Response~(session*cue)+(MeanFD)+(session*cue|subjectID)’).*

To test whether these responses were correlated with individual differences in self-reported craving, we calculated the average contrast value (food>flowers and social>flowers) in the anatomically defined SN/VTA for each participant. We used two different approaches to measure participants’ self-reported craving. First, we calculated the mean craving rating participants reported on each trial during the CIC task in the scanner (*Craving_CIC*). This measure was exactly comparable across sessions, and simultaneous with the fMRI data acquisition. Because the data for food craving were truncated at the upper limit of 10 (see results figure 3), we also calculated a truncated regression model (using the truncreg package in R) using the truncated variable as the explanatory variable in the regression model^88,89^ in addition to the standard correlation. Second, we took the craving reported by participants on the final online questionnaire, completed after 10 hours of fasting or isolation (*Craving_Q*). On this measure participant only reported craving for the deprived need (food when fasting, social contact when isolated), but this measure provides the most direct measure of the effect of deprivation because it can be compared to the self-report at the beginning of each session. In addition, the *Craving_Q* ratings were completed on a finer scale (0-100 instead of 0-10) and participants had no time restriction when filling out the questionnaires (while during the task, participants had 5s to complete the scales). For these reasons, we include both types of craving ratings and report results as significant at α<0.025. We measured correlations between self-reported craving and neural responses for each deprivation session. Because we specifically predicted a positive correlation between craving for the deprived target and response magnitude in SN/VTA, these correlations were tested one-tailed. Data for the analyses was extracted using FSL’s ‘fslmeants’ utility and subsequent univariate and correlation analyses were conducted in Matlab 2019b, RStudio (1.1.423) and SPSS 26.

#### Multivariate Analyses

We next used multivoxel pattern analysis (MVPA) to determine whether the multivariate spatial pattern of activity in SN/VTA is shared for food and social craving. From the GLM, we extracted the beta values (i.e., amplitude of the fit hemodynamic response function) of the response to each condition (food cues, social cues, flower cues) for each block in each run (3 blocks per condition per run * 6 runs) in each session (baseline, fasting, isolation) resulting in 162 beta values for each voxel. Responses were extracted from all voxels in the anatomically identified SN/VTA in each participant (i.e. no additional feature selection was applied). All multivariate analyses were conducted with the PyMVPA^90^ toolbox in Python (http://www.pymvpa.org) and Matlab2019b. While smoothing has been shown to not substantially affect information in multivariate data^91–93^, because of the small size of the SN/VTA, we also ran the classification analyses on unsmoothed data and find that we were not able to decode stimulus or motivational state across sessions in the unsmoothed data (see Supplementary Materials).

First, as a proof of concept, we tested if we could classify food cues from control cues from the SN/VTA within the food fasting session (see Supplementary Materials). Then, we trained a linear SVM classifier, training using all 36 betas (2 cues * 3 blocks * 6 runs) of the food fasting session to discriminate patterns of responses to food vs flower cues (see Supplementary Materials for within-session decoding accuracies for food cues). We tested the generalization of the classifier to responses to social vs flower cues in the isolation session (i.e., different category of cue but similar motivational state), producing an accuracy score (correct classification for 36 betas). If social craving and food craving share a neural basis, we predicted that a classifier trained on food_craved vs control cues would successfully (above chance) classify social_craved vs control cues. We also tested the generalization of this classifier to food vs flower cues, in the isolation and baseline sessions (i.e., the same categories of cues as the training, but different motivational state), and to social vs flower cues at baseline (i.e., different category and different motivational state, as a control analysis). In order to obtain confidence intervals of the mean in the data samples we used bootstrapping. We generated 1000 datasets randomly by sampling with replacement from the classification accuracies across participants using Matlab’s bootci function.

For hypothesis testing on the group level, we used a permutation analysis following the methods in Stelzer et al. 2013^94^. This non-parametric approach does not depend on assumptions about the distribution of classification accuracies^94,95^. To generate a null distribution from the data, we followed the steps described in Stelzer et al., which we summarize below and visualize in figure 4. We shuffled the condition labels randomly during training within each run, and then tested the prediction accuracies for each cross-classification on the test data. We performed this permutation analysis 100 times per participant (thus creating 100 random permutations) resulting in 100 accuracy values per participant, for each testing data set. To create a null distribution on the group level, we then randomly drew one of the 100 accuracy values for each participant, and calculated a mean across participants. This procedure was repeated 10^5^ times for each testing data set, creating the null distributions for each data set (see figure 4 for the histograms showing the null distributions). We calculated the probability p of obtaining a mean accuracy value in the null distributions that is equal to or higher than the true mean from the analyses. Following Stelzer et al., we rejected the null hypothesis of no group-level decoding if p < 0.001 which corresponds to a low probability if it were retrieved by chance.

Finally, we used representational similarity analysis to test which pattern of activity is more similar to ‘food_craved’: social_craved or food_noncraved. We predicted that the presence of a craved object should be more important for SN/VTA activity than the cue category, so we predicted that the pattern of food_craved responses will be *more similar* to the pattern of response to social_craved cues, compared to when these cues are presented in different motivational states (social_craved x food_noncraved) or even the same cue in different motivational states (food_craved x food_noncraved). We averaged responses to each cue across all 6 runs per session to obtain one mean beta value for each voxel. Then we calculated the dissimilarity (1 – Pearson’s correlation, fisher-transformed) between each average response pattern. Then we compared the dissimilarity between food_craved x social_craved (i.e., food-fasted and social-isolated; different stimuli but similar motivational state) to the dissimilarity between food_noncraved x social_craved (i.e., food-baseline and social-isolate; maximally distant (different stimuli, different motivational state and different session – the noise ceiling) and to the dissimilarity between food_noncraved x food_craved (i.e., food-isolation and food-fasted; same stimuli but different motivational state (we used isolation instead of baseline as it represents the satiated control condition to fasting)); both comparisons used pair-wise t-tests. Data distribution was assumed to be normal but this was not formally tested.

#### Correlations univariate fMRI measures and behavioral data

In follow-up analyses, we tested whether individual differences in social network size predict the magnitude of self-reported and neural measures of social craving. We used the UCLA loneliness score obtained from each participant as an indicator of their chronic loneliness. We correlated these two measures with mean values extracted from the midbrain and striatum ROIs from the contrast social_craved>control.

### Additional analyses

#### SN/VTA: Multivariate decoding of food cues

As a proof of concept, we tested if we could discriminate fMRI patterns between food and control cues in the fasting session in the SN/VTA. For each participant, we partitioned the data into six independent folds (6 runs), and iteratively trained a linear support vector machine (SVM) classifier on 5 runs (i.e. 15 beta estimates per condition) and tested on the left-out run (3 beta estimates per condition). We then averaged the classification accuracy across runs to yield a single estimate for each participant. This within-session classification tested whether we would be able to decode the cue type (food vs. control) from multivariate patterns within the SN/VTA. We tested whether the accuracy was significantly above chance using bootstrapping. We first generated a null-distribution of the data by shifting the mean to be 0.5 (by calculating dataset – mean(dataset) + 0.5) and then generated 1000 datasets randomly by sampling with replacement from the classification accuracies across subjects using Matlab’s bootstrp function. We then calculated a mean for each of the 1000 simulated datasets from the null distribution and compared it to our observed data average. We calculated the probability p of obtaining a mean value across the 1000 datasets that is equal or higher than the actual mean from the original dataset and rejected the null hypothesis of no group-level decoding if p < 0.05. For obtaining confidence intervals of the mean in the data samples, we generated 1000 datasets randomly by sampling with replacement from the classification accuracies across subjects using Matlab’s bootci function. We found above chance (50%) decoding of food cues from flower images after fasting (mean accuracy=0.5556; p<0.001, 95% CI=0.527-0.585).

#### SN/VTA: Cross-classification of food and social cues - unsmoothed data

We ran the same analysis as reported in the main manuscript on data for which we did not perform any smoothing during preprocessing in order to test whether smoothing has substantial effects on our results. First, we tested how smoothing affected the accuracy to discriminate fMRI patterns between food and control cues in the fasting session in the SN/VTA using the same method as described above (“SN/VTA: Multivariate decoding of food cues) on the unsmoothed data. Here, we find that omitting smoothing showed slightly lower mean accuracy across subjects (mean accuracy=0.542) but still resulted in successful above chance (50%) decoding of food cues from flower images after fasting (p=0.002, 95% CI=0.513-0.570). Then, as in the original analysis, we trained a classifier on the pattern of activity in the SN/VTA after fasting to food cues and flower images and tested whether it would successfully decode food and social cues from control cues in the other two sessions (baseline and isolation). Again, we generated null distributions using the permutation analysis described in the main text (following Stelzer et al. 2013^94^). In the unsmoothed data, we find no above chance decoding (α=0.001) of social cues from flowers after isolation (mean accuracy=0.521, bootstrapped CI=[0.503,0.542], p=0.041) or after baseline (mean accuracy=0.526, bootstrapped CI=[0.501,0.546], p=0.021). In addition, in unsmoothed data, the classifier was also not able to decode food cues from flowers after isolation and baseline (isolation: mean accuracy=0.522, bootstrapped CI=[0.500,0.544], p=0.021; baseline: mean accuracy=0.525, bootstrapped CI=[0.501,0.557], p=0.022).

#### Exploratory whole brain group analyses

As a complementary analysis, we assessed main effects and interactions of cue and session on whole brain activity. We set up a flexible factorial model with the factors subject, session (baseline, fasting, isolation) and cue (food, social). We entered the first-level contrasts food > control and social > control for each session into the group-level analysis. Analyses were implemented in SPM12b.

##### Main effects: Cue

We first assessed whole mean brain activity for the contrasts: food > control and social > control across all experimental sessions (baseline, fasting, isolation) as a manipulation check to assess which regions were activated in response to food (vs. control) and social (vs. control) cues. We corrected for multiple comparisons using whole brain family-wise error correction. Table S21 and Figure S7 **s**how the results from this analysis.

##### Interaction effects: Cue * deprivation

We then assessed effects of isolation on brain activity in response to social cues and effects of fasting on brain activity in response to food cues in the whole brain. To do this, we calculated two contrasts: 1) food: fasting > isolation and 2) social: isolation > fasting. These analyses were exploratory, and statistical inference was performed using a threshold of p < 0.05 corrected for multiple comparisons over the whole brain, using the Gaussian random fields approach at cluster-level with a voxel-level intensity threshold of p < 0.001^96^. We note that this method of correction for multiple comparisons is likely biased towards false positives^97,98^. We decided to share these lenient analyses (Tables S22 / S23 and Figure S8) to be more complete and for potential future meta-analyses.

#### Midbrain localizer: group analysis

As described above, we used a functional localizer to select the most active voxels for reward anticipation and novelty within the midbrain to functionally localize the SN/VTA in each participant individually. To explore the localization of midbrain responses to the localizer task in more detail, we implemented a group random effects analysis entering the first level contrast reward > non-reward from each participant (see Figure S9).

Data analysis was not performed blind to the experimental conditions.

## Data and materials availability

All de-identified neuroimaging data are publicly available on OpenNeuro.org / DOI: 10.18112/openneuro.ds003242.v1.0.0. Summary fMRI and behavioral data are publicly available on the Open Science Framework (OSF) / DOI: DOI 10.17605/OSF.IO/F9CRU. Stimuli for the tasks were taken from the open image sharing website www.pexels.com.

## Code availability

Analysis code, code to generate the figures is publicly available on OSF / DOI: DOI 10.17605/OSF.IO/F9CRU. Code to run the tasks with example stimuli is also publicly available on OSF / DOI: 10.17605/OSF.IO/CF2RT.

## Acknowledgements

This research was carried out at the Athinoula A. Martinos Imaging Center at the McGovern Institute for Brain Research at MIT. The authors thank Katherine Sottilare, Molly Humphreys, James Huettig, Rania Ezzo, Javier Weddington, Joachim Kennedy, Michelle Hung and Isabel Nichoson for help with data collection, and Diana Tamir, Judith Mildner, Hilary Richardson, Shari Liu, Emrah Duzel, Nancy Kanwisher and Daniel Nettle for advice and discussion. We also thank Akhil Gupta for his help making the fMRI dataset publicly available. We gratefully acknowledge support of this project by a SFARI Explorer Grant from the Simons Foundation (#597310 to R.S.), a MINT grant from the McGovern Institute (#1496911to R.S.), an NIH Pioneer Award (#DP1-AT009925 to K.T.), a Max Kade Foundation fellowship (to L.T.), an Erwin Schroedinger Fellowship by the Austrian Science Fund (# J4326-B25 to L.T.) and an NIH shared instrumentation grant (#1S10OD021569-01). RS participated in the Center for Brains, Minds and Machines (CBMM), funded by a NSF STC award (CCF-1231216).

## Author contributions

L.T. and R.S. designed the study with input from K.T. and G.M.; A.T. provided support in optimizing the scanning parameters and during fMRI data collection. L.T. collected the data with support from K.W.; L.T. analyzed the data with support from T.T. and R.S.; L.T. and R.S. wrote the manuscript and all authors provided feedback on the final version.

## Competing interests

The authors declare no competing interests.

## Notes

https://osf.io/cwg9e/

### Competing Interest Statement

The authors have declared no competing interest.

### Summary of Updates

Revised manuscript version following peer review

