## Supplemental file for "Acute social isolation evokes midbrain craving responses similar to hunger"

### Supplementary Figures

##### Craving ratings all sessions

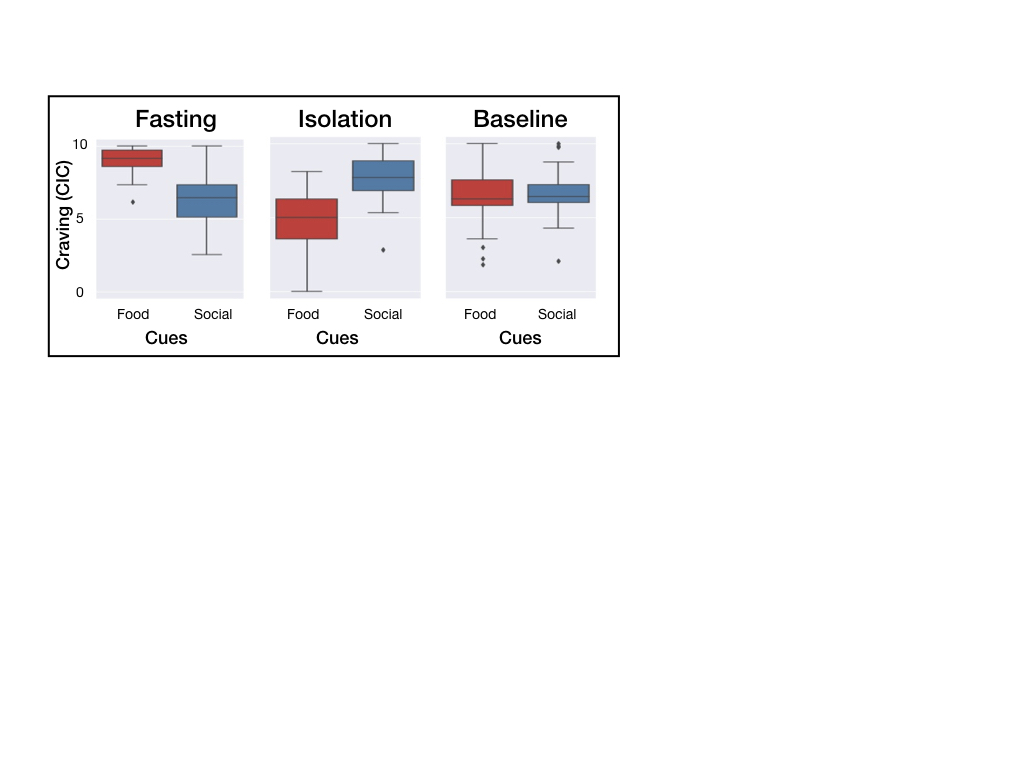

Supplementary Figure 1. Craving ratings (n = 40) during the cue induced craving task in response to food cues and social cues for each session. The boxplots in indicate the median (dark center line), the interquartile range (IQR; box) and the 1.5 IQR minima and maxima (whiskers). Datapoints outside the whiskers are shown as individual data points.

##### SN/VTA: Full data for all sessions and all cues

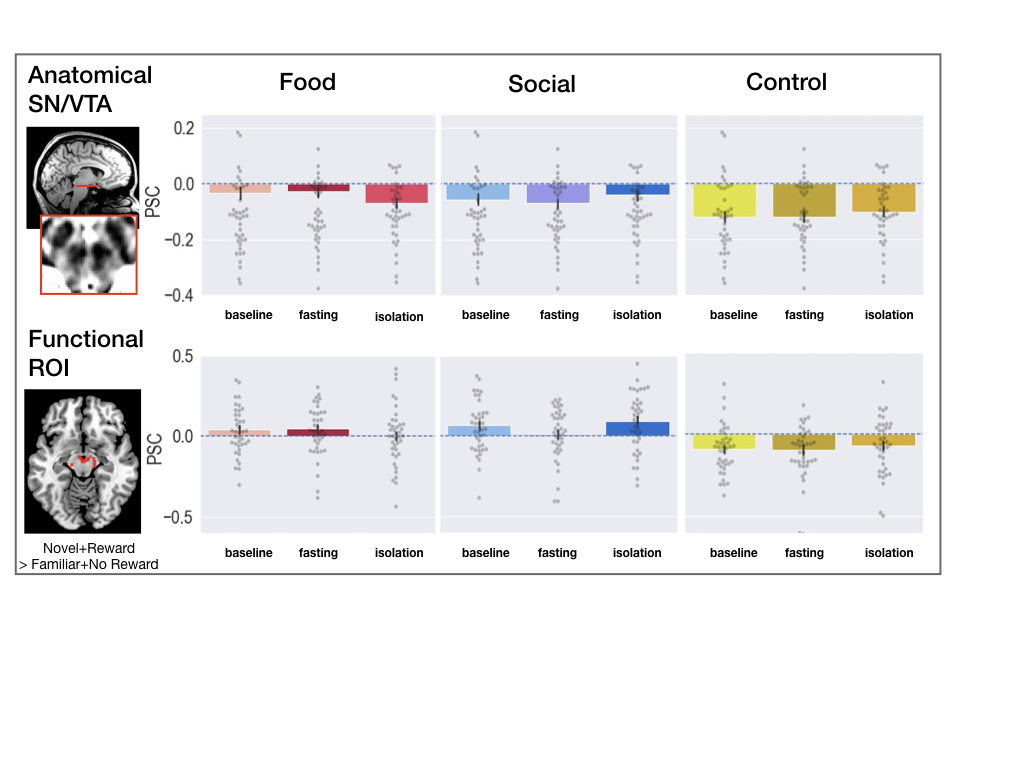

Supplementary Figure 2. Full data (n = 40): Univariate activity in the SN/VTA (upper row)^[1]^ and the midbrain functional ROI (lower row), for all cues and all sessions. The bar plots depict the mean beta values for food, social and control cues. The grey dots indicate individual data points and the error bars indicate standard errors of the mean. The dashed blue horizontal line indicates zero.

##### NAcc – functional ROI: Full data for all sessions and all cues

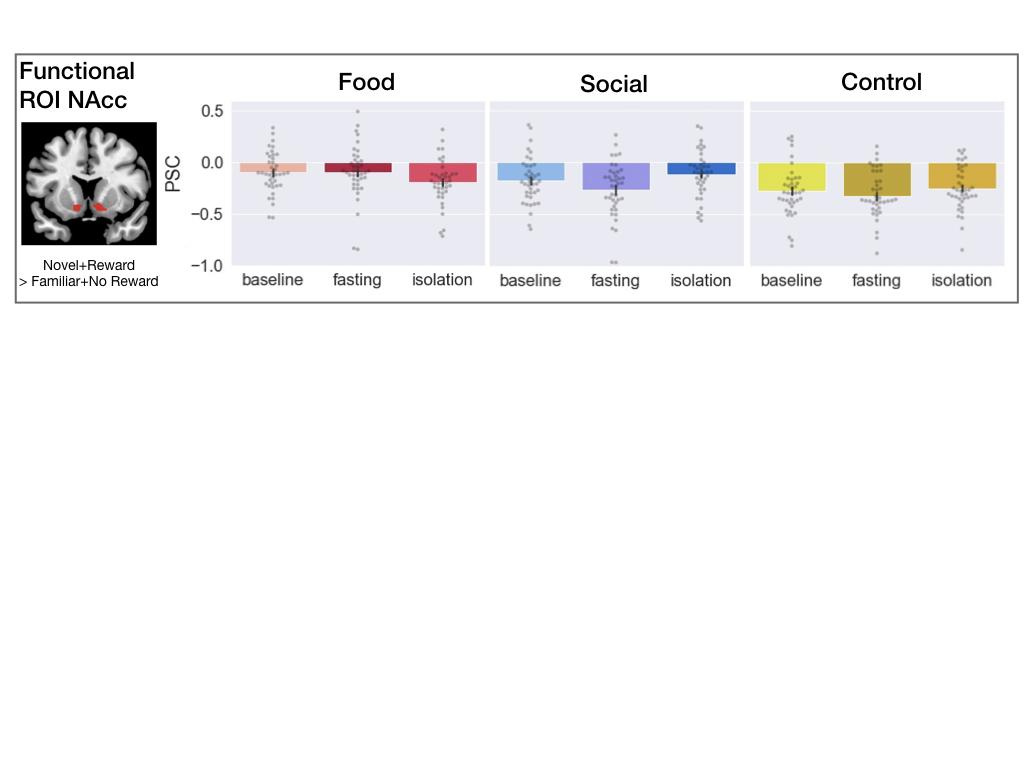

Supplementary Figure 3. Univariate activity in the NAcc functional ROI for all cues and all sessions (n = 40). The bar plots depict the mean beta values for food, social and control cues. The dots indicate individual data points and the error bars indicate standard errors of the mean.

##### Group level whole brain: session * cue interaction

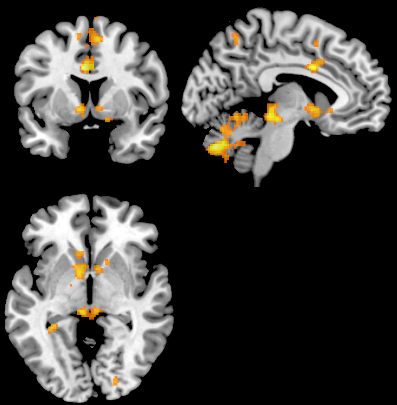

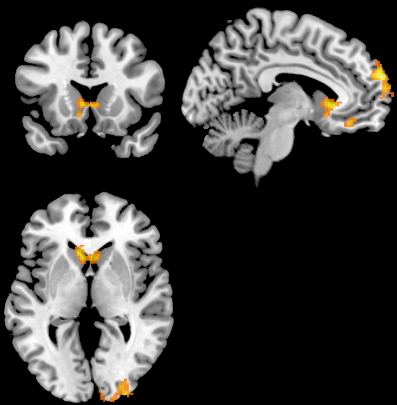

**Food: Fasting>Isolation**

**Social: Isolation>Fasting**

Supplementary Figure 4. Univariate group-level activity cluster-level corrected over the whole brain (n = 40). Left: contrast food>control: fasting > isolation; Right: contrast social>control: isolation > fasting. Tables 21 (food > control) and 22 (social > control) show the results for this analysis.

##### Correlations between craving ratings

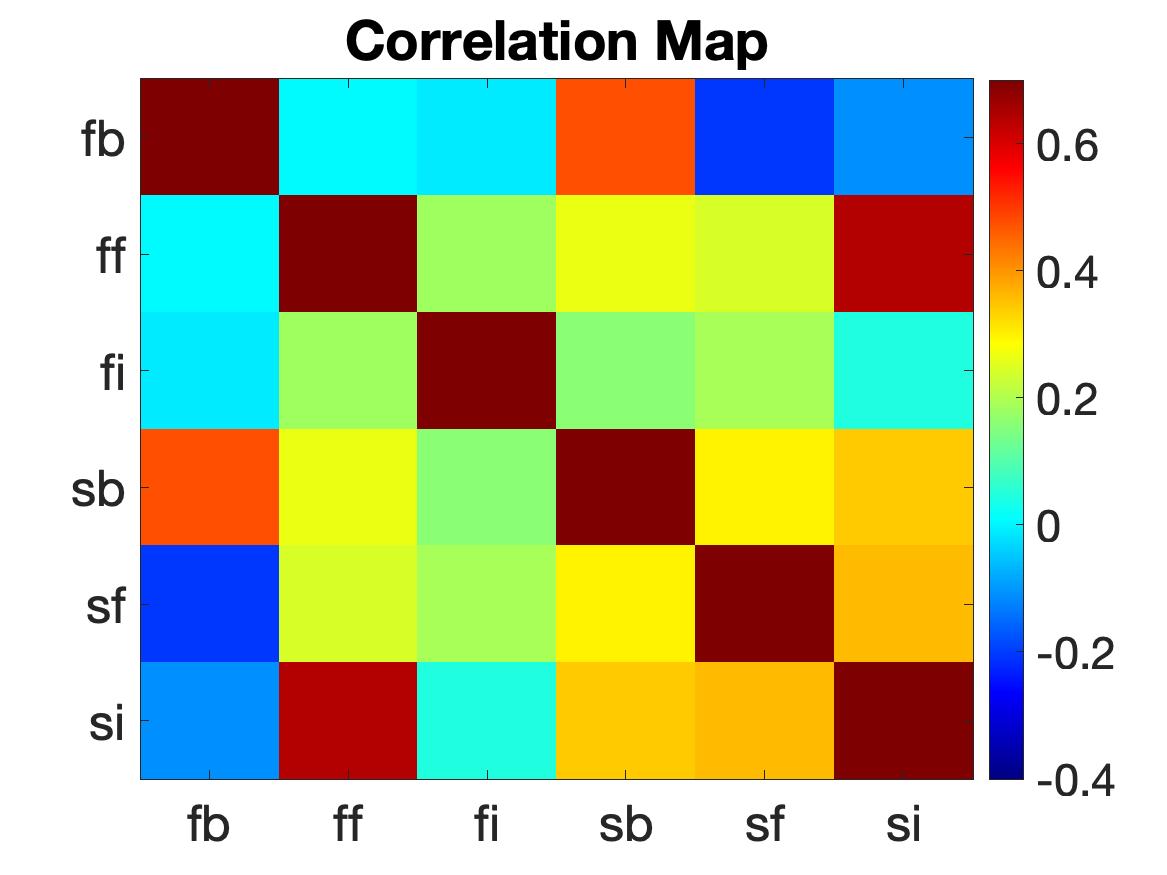

Supplementary Figure 5. Correlations between craving ratings during the cue induced craving task for each session (n = 40). fb = food craving, baseline session; ff = food craving, fasting session; fi = food craving, isolation session; sb = social craving, baseline session; sf = social craving, fasting session; si = social craving, isolation session.

##### Group level whole brain: main of effects of cue

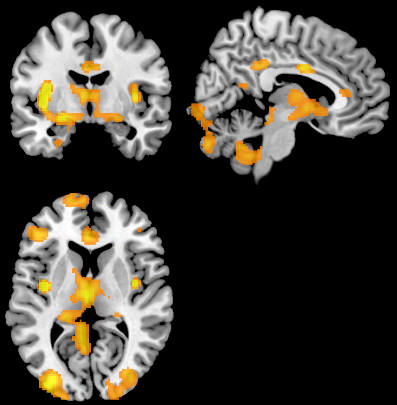

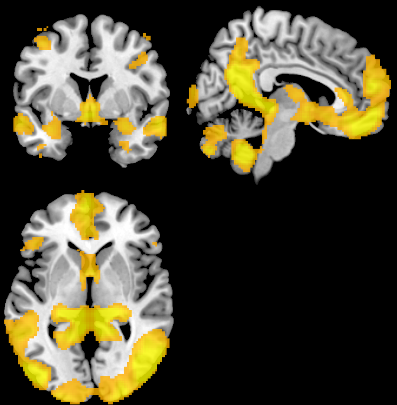

**Food>Control**

**Social>Control**

Supplementary Figure 6. Univariate whole brain group-level activity family-wise error corrected at the voxel level (n = 40) : Left: contrast food>control (mean across all sessions: baseline, fasting, isolation). Right: social>control (mean across all sessions). Table 23 shows the results for this analysis.

##### Midbrain localizer: group analysis

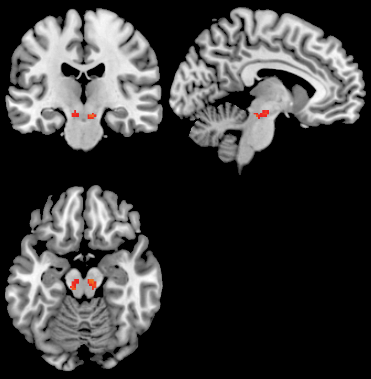

**Localizer: Reward>Nonreward**

Supplementary Figure 7. Univariate group-level activity in the midbrain localizer task within the midbrain for the contrast reward > nonreward (n = 40). All voxels with p<0.001 within the midbrain are displayed (no correction for multiple comparisons).

##### Comparison anatomical and functional SN/VTA ROIs

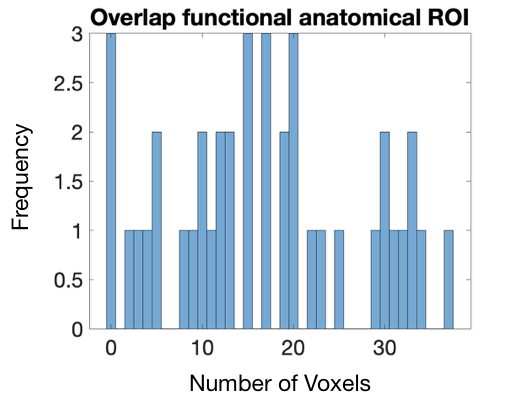

Supplementary Figure 8. Number of overlapping voxels between the functional and anatomical midbrain ROI (n = 40). The overlap ranged between 0-30 voxels out of a possible 100 voxels.

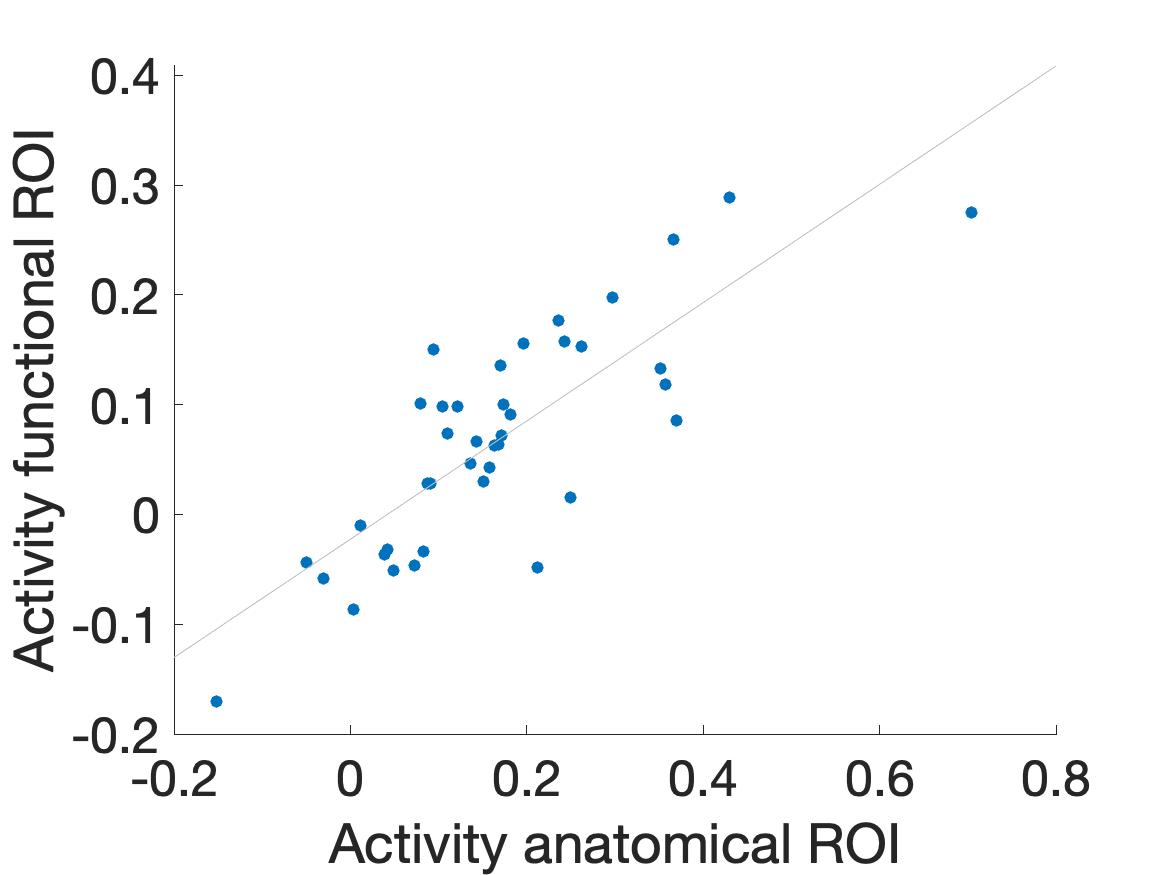

Supplementary Figure 9. Correlation between activity in functional and anatomical ROIs across subjects (n = 40).

### Supplementary Tables

##### Results: SN/VTA

###### Fasting vs Isolation contrast, anatomical SN/VTA

| Predictors | Estimates | t | p |
| --- | --- | --- | --- |
| Session: Isolation > Fasting | 0.009 | 1.2 | 0.230 |
| Cue: Food | 0.08 | 7.0 | 3.13e-11 |
| Cue: Social | 0.06 | 5.2 | 4.28e-7 |
| Interaction: session – cue (food) | -0.03 | -4.0 | 0.0005 |
| Interaction: session – cue (social) | 0.006 | 0.6 | 0.535 |

Supplementary Table 1. *Output mixed effects model: Fasting vs. Isolation contrast, SN/VTA anatomical ROI.* Results are shown for a mixed effects regressions (using Matlab 2019b’s *fitlme* function) to estimate the fixed effects of cue, deprivation session, and their interaction on response magnitude in the ROI, controlling for each session’s average framewise displacement (i.e. head motion), with participant included as a random effect with both random intercepts and random slopes. The reported p-values (two-tailed) are not corrected for multiple comparisons.

###### Fasting vs Isolation contrast, midbrain functional ROI

| Predictors | Estimates | t | p |
| --- | --- | --- | --- |
| Session: Isolation > Fasting | 0.006 | 0.36 | 0.720 |
| Cue: Food | 0.11 | 6.7 | 1.60e-10 |
| Cue: Social | 0.13 | 7.2 | 1.04e-11 |
| Interaction: session – cue (food) | -0.03 | -3.3 | 0.001 |
| Interaction: session – cue (social) | 0.03 | 2.5 | 0.015 |

Supplementary Table 2. *Output mixed effects model: Fasting vs. Isolation contrast, midbrain functional ROI.* Results are shown for a mixed effects regressions (using Matlab 2019b’s *fitlme* function) to estimate the fixed effects of cue, deprivation session, and their interaction on response magnitude in the ROI, controlling for each session’s average framewise displacement (i.e. head motion), with participant included as a random effect with both random intercepts and random slopes. The reported p-values (two-tailed) are not corrected for multiple comparisons.

###### Fasting and Isolation compared to baseline session, anatomical SN/VTA

| Predictors | Estimates | t | p |
| --- | --- | --- | --- |
| Session: Fasting | 0.01 | 0.3 | 0.795 |
| Session: Isolation | 0.01 | 0.7 | 0.496 |
| Cue: Food | 0.09 | 4.0 | 8.06e-5 |
| Cue: Social | 0.06 | 3.0 | 0.003 |
| Interaction: session (fasting) – cue (food) | 0.005 | 0.2 | 0.836 |
| Interaction: session (isolation) – cue (food) | -0.05 | -2.1 | 0.038 |
| Interaction: session (fasting) – cue (social) | -0.01 | -0.6 | 0.576 |
| Interaction: session (isolation) – cue (social) | -0.002 | -0.1 | 0.941 |

Supplementary Table 3. *Output mixed effects model: Fasting and Isolation compared to Baseline, SN/VTA anatomical ROI.* Results are shown for a mixed effects regressions (using Matlab 2019b’s *fitlme* function) to estimate the fixed effects of cue, deprivation session, and their interaction on response magnitude in the ROI, controlling for each session’s average framewise displacement (i.e. head motion), with participant included as a random effect with both random intercepts and random slopes. The reported p-values (two-tailed) are not corrected for multiple comparisons.

###### Fasting and Isolation compared to baseline session, midbrain functional ROI

| Predictors | Estimates | t | p |
| --- | --- | --- | --- |
| Session: Fasting | 0.005 | 0.2 | 0.857 |
| Session: Isolation | 0.02 | 0.6 | 0.531 |
| Cue: Food | 0.14 | 5.2 | 2.67e-7 |
| Cue: Social | 0.16 | 5.8 | 1.14e-8 |
| Interaction: session (fasting) – cue (food) | 0.004 | 0.14 | 0.900 |
| Interaction: session (isolation) – cue (food) | -0.06 | -2.4 | 0.018 |
| Interaction: session (fasting) – cue (social) | -0.05 | -1.8 | 0.070 |
| Interaction: session (isolation) – cue (social) | 0.004 | 0.14 | 0.900 |

Supplementary Table 4*. Output mixed effects model: Fasting and Isolation compared to Baseline, midbrain functional ROI.* Results are shown for a mixed effects regressions (using Matlab 2019b’s *fitlme* function) to estimate the fixed effects of cue, deprivation session, and their interaction on response magnitude in the ROI, controlling for each session’s average framewise displacement (i.e. head motion), with participant included as a random effect with both random intercepts and random slopes. The reported p-values (two-tailed) are not corrected for multiple comparisons.

##### Putamen

###### Fasting vs Isolation contrast

| Predictors | Estimates | t | p |
| --- | --- | --- | --- |
| Session: Isolation > Fasting | 0.003 | 0.3 | 0.751 |
| Cue: Food | 0.024 | 1.8 | 0.077 |
| Cue: Social | -0.006 | -0.4 | 0.673 |
| Interaction: session * cue (food) | -0.033 | -3.2 | 0.002 |
| Interaction: session * cue (social) | 0.016 | 1.4 | 0.173 |

Supplementary Table 5. *Output mixed effects model: Fasting vs. Isolation contrast, Putamen ROI.* Results are shown for a mixed effects regressions (using Matlab 2019b’s *fitlme* function) to estimate the fixed effects of cue, deprivation session, and their interaction on response magnitude in the ROI, controlling for each session’s average framewise displacement (i.e. head motion), with participant included as a random effect with both random intercepts and random slopes. The reported p-values (two-tailed) are not corrected for multiple comparisons.

###### Comparison to baseline session

| Predictors | Estimates | t | p |
| --- | --- | --- | --- |
| Session: Fasting | 0.004 | 0.1 | 0.886 |
| Session: Isolation | 0.012 | 0.4 | 0.676 |
| Cue: Food | 0.047 | 2.2 | 0.031 |
| Cue: Social | 0.011 | 0.6 | 0.535 |
| Interaction: session (fasting) – cue (food) | 0.010 | 0.3 | 0.742 |
| Interaction: session (isolation) – cue (food) | -0.060 | -2.5 | 0.012 |
| Interaction: session (fasting) – cue (social) | -0.033 | -1.2 | 0.222 |
| Interaction: session (isolation) – cue (social) | -0.002 | -0.1 | 0.935 |

Supplementary Table 6. *Output mixed effects model: Fasting and Isolation compared to Baseline, Putamen ROI.* Results are shown for a mixed effects regressions (using Matlab 2019b’s *fitlme* function) to estimate the fixed effects of cue, deprivation session, and their interaction on response magnitude in the ROI, controlling for each session’s average framewise displacement (i.e. head motion), with participant included as a random effect with both random intercepts and random slopes. The reported p-values (two-tailed) are not corrected for multiple comparisons.

##### NAcc

###### Fasting vs Isolation contrast

| Predictors | Estimates | t | p |
| --- | --- | --- | --- |
| Session: Isolation > Fasting | 0.033 | 1.9 | 0.060 |
| Cue: Food | 0.128 | 6.4 | 1.09e-9 |
| Cue: Social | 0.114 | 5.2 | 3.70e-7 |
| Interaction: session * cue (food) | -0.080 | -4.7 | 4.51e-6 |
| Interaction: session * cue (social) | 0.038 | 2.1 | 0.040 |

Supplementary Table 7. *Output mixed effects model: Fasting vs. Isolation contrast, NAcc ROI.* Results are shown for a mixed effects regressions (using Matlab 2019b’s *fitlme* function) to estimate the fixed effects of cue, deprivation session, and their interaction on response magnitude in the ROI, controlling for each session’s average framewise displacement (i.e. head motion), with participant included as a random effect with both random intercepts and random slopes. The reported p-values (two-tailed) are not corrected for multiple comparisons.

###### Comparison to baseline session

| Predictors | Estimates | t | p |
| --- | --- | --- | --- |
| Session: Fasting | -0.051 | -1.2 | 0.241 |
| Session: Isolation | 0.020 | 0.5 | 0.643 |
| Cue: Food | 0.161 | 5.7 | 2.87e-8 |
| Cue: Social | 0.104 | 3.3 | 0.001 |
| Interaction: session (fasting) – cue (food) | 0.048 | 1.1 | 0.260 |
| Interaction: session (isolation) – cue (food) | -0.113 | -3.0 | 0.004 |
| Interaction: session (fasting) – cue (social) | -0.028 | -0.8 | 0.455 |
| Interaction: session (isolation) – cue (social) | 0.050 | 1.3 | 0.192 |

Supplementary Table 8. *Output mixed effects model: Fasting and Isolation compared to Baseline, NAcc ROI.* Results are shown for a mixed effects regressions (using Matlab 2019b’s *fitlme* function) to estimate the fixed effects of cue, deprivation session, and their interaction on response magnitude in the ROI, controlling for each session’s average framewise displacement (i.e. head motion), with participant included as a random effect with both random intercepts and random slopes. The reported p-values (two-tailed) are not corrected for multiple comparisons.

##### NAcc – exploratory functional ROI

###### Fasting vs Isolation contrast

| Predictors | Estimates | t | p |
| --- | --- | --- | --- |
| Session: Isolation > Fasting | 0.030 | 1.6 | 0.122 |
| Cue: Food | 0.142 | 6.1 | 4.70e-9 |
| Cue: Social | 0.100 | 4.2 | 3.40e-5 |
| Interaction: session * cue (food) | -0.090 | -4.3 | 2.23e-5 |
| Interaction: session * cue (social) | 0.038 | 1.9 | 0.061 |

Supplementary Table 9. *Output mixed effects model: Fasting vs. Isolation contrast, NAcc functional ROI.* Results are shown for a mixed effects regressions (using Matlab 2019b’s *fitlme* function) to estimate the fixed effects of cue, deprivation session, and their interaction on response magnitude in the ROI, controlling for each session’s average framewise displacement (i.e. head motion), with participant included as a random effect with both random intercepts and random slopes. The reported p-values (two-tailed) are not corrected for multiple comparisons.

###### Comparison to baseline session

| Predictors | Estimates | t | p |
| --- | --- | --- | --- |
| Session: Fasting | -0.042 | -0.9 | 0.384 |
| Session: Isolation | 0.020 | 0.4 | 0.657 |
| Cue: Food | 0.176 | 6.1 | 2.71e-9 |
| Cue: Social | 0.100 | 3.3 | 0.001 |
| Interaction: session (fasting) – cue (food) | 0.052 | 1.1 | 0.292 |
| Interaction: session (isolation) – cue (food) | -0.121 | -3.0 | 0.004 |
| Interaction: session (fasting) – cue (social) | -0.040 | -1.0 | 0.315 |
| Interaction: session (isolation) – cue (social) | 0.040 | 1.0 | 0.322 |

Supplementary Table 10. *Output mixed effects model: Fasting and Isolation compared to Baseline, NAcc functional ROI.* Results are shown for a mixed effects regressions (using Matlab 2019b’s *fitlme* function) to estimate the fixed effects of cue, deprivation session, and their interaction on response magnitude in the ROI, controlling for each session’s average framewise displacement (i.e. head motion), with participant included as a random effect with both random intercepts and random slopes. The reported p-values (two-tailed) are not corrected for multiple comparisons.

##### Caudate

###### Fasting vs Isolation contrast

| Predictors | Estimates | t | p |
| --- | --- | --- | --- |
| Session: Isolation > Fasting | -0.003 | -0.2 | 0.849 |
| Cue: Food | 0.058 | 4.0 | 8.49e-5 |
| Cue: Social | 0.023 | 1.3 | 0.181 |
| Interaction: session * cue (food) | -0.025 | -1.9 | 0.061 |
| Interaction: session * cue (social) | 0.035 | 3.0 | 0.003 |

Supplementary Table 11. *Output mixed effects model: Fasting vs. Isolation contrast, Caudate ROI.* Results are shown for a mixed effects regressions (using Matlab 2019b’s *fitlme* function) to estimate the fixed effects of cue, deprivation session, and their interaction on response magnitude in the ROI, controlling for each session’s average framewise displacement (i.e. head motion), with participant included as a random effect with both random intercepts and random slopes. The reported p-values (two-tailed) are not corrected for multiple comparisons.

###### Comparison to baseline session

| Predictors | Estimates | t | p |
| --- | --- | --- | --- |
| Session: Fasting | 0.011 | 0.3 | 0.740 |
| Session: Isolation | 0.006 | 0.2 | 0.860 |
| Cue: Food | 0.077 | 4.0 | 8.12e-5 |
| Cue: Social | 0.010 | 0.5 | 0.614 |
| Interaction: session (fasting) – cue (food) | 0.007 | 0.2 | 0.832 |
| Interaction: session (isolation) – cue (food) | -0.044 | -2.0 | 0.053 |
| Interaction: session (fasting) – cue (social) | -0.022 | -0.8 | 0.440 |
| Interaction: session (isolation) – cue (social) | 0.048 | 2.2 | 0.033 |

Supplementary Table 12. *Output mixed effects model: Fasting and Isolation compared to Baseline, Caudate ROI.* Results are shown for a mixed effects regressions (using Matlab 2019b’s *fitlme* function) to estimate the fixed effects of cue, deprivation session, and their interaction on response magnitude in the ROI, controlling for each session’s average framewise displacement (i.e. head motion), with participant included as a random effect with both random intercepts and random slopes. The reported p-values (two-tailed) are not corrected for multiple comparisons.

##### Orbitofrontal cortex (OFC)

###### Fasting vs Isolation contrast

| Predictors | Estimates | t | p |
| --- | --- | --- | --- |
| Session: Isolation > Fasting | -0.006 | -0.2 | 0.857 |
| Cue: Food | 0.249 | 5.5 | 1.26e-7 |
| Cue: Social | 0.968 | 12.5 | 1.09e-27 |
| Interaction: session * cue (food) | -0.033 | -1.0 | 0.337 |
| Interaction: session * cue (social) | 0.112 | 2.5 | 0.0121 |

Supplementary Table 13. *Output mixed effects model: Fasting vs. Isolation contrast, OFC ROI.* Results are shown for a mixed effects regressions (using Matlab 2019b’s *fitlme* function) to estimate the fixed effects of cue, deprivation session, and their interaction on response magnitude in the ROI, controlling for each session’s average framewise displacement (i.e. head motion), with participant included as a random effect with both random intercepts and random slopes. The reported p-values (two-tailed) are not corrected for multiple comparisons.

###### Comparison to baseline session

| Predictors | Estimates | t | p |
| --- | --- | --- | --- |
| Session: Fasting | 0.079 | 1.0 | 0.311 |
| Session: Isolation | 0.078 | 1.0 | 0.330 |
| Cue: Food | 0.288 | 4.6 | 6.35e-6 |
| Cue: Social | 0.897 | 9.5 | 2.56e-19 |
| Interaction: session (fasting) – cue (food) | -0.006 | -0.1 | 0.926 |
| Interaction: session (isolation) – cue (food) | -0.072 | -0.8 | 0.416 |
| Interaction: session (fasting) – cue (social) | -0.041 | -0.5 | 0.601 |
| Interaction: session (isolation) – cue (social) | 0.184 | 2.1 | 0.039 |

Supplementary Table 14. *Output mixed effects model: Fasting and Isolation compared to Baseline, OFC ROI.* Results are shown for a mixed effects regressions (using Matlab 2019b’s *fitlme* function) to estimate the fixed effects of cue, deprivation session, and their interaction on response magnitude in the ROI, controlling for each session’s average framewise displacement (i.e. head motion), with participant included as a random effect with both random intercepts and random slopes. The reported p-values (two-tailed) are not corrected for multiple comparisons.

##### Amygdala

###### Fasting vs Isolation contrast

| Predictors | Estimates | t | p |
| --- | --- | --- | --- |
| Session: Isolation > Fasting | 0.026 | 1.4 | 0.160 |
| Cue: Food | 0.145 | 6.4 | 1.04e-9 |
| Cue: Social | 0.259 | 9.2 | 1.78e-17 |
| Interaction: session * cue (food) | -0.035 | -1.9 | 0.055 |
| Interaction: session * cue (social) | 0.019 | 1.0 | 0.334 |

Supplementary Table 15. *Output mixed effects model: Fasting vs. Isolation contrast, Amygdala ROI.* Results are shown for a mixed effects regressions (using Matlab 2019b’s *fitlme* function) to estimate the fixed effects of cue, deprivation session, and their interaction on response magnitude in the ROI, controlling for each session’s average framewise displacement (i.e. head motion), with participant included as a random effect with both random intercepts and random slopes. The reported p-values (two-tailed) are not corrected for multiple comparisons.

###### Comparison to baseline session

| Predictors | Estimates | t | p |
| --- | --- | --- | --- |
| Session: Fasting | -0.022 | -0.6 | 0.548 |
| Session: Isolation | 0.029 | 0.7 | 0.463 |
| Cue: Food | 0.142 | 5.3 | 1.89e-7 |
| Cue: Social | 0.275 | 6.6 | 1.78e-10 |
| Interaction: session (fasting) – cue (food) | 0.037 | 0.7 | 0.300 |
| Interaction: session (isolation) – cue (food) | -0.032 | -0.8 | 0.404 |
| Interaction: session (fasting) – cue (social) | -0.035 | -0.9 | 0.391 |
| Interaction: session (isolation) – cue (social) | 0.003 | 0.1 | 0.941 |

Supplementary Table 16. *Output mixed effects model: Fasting and Isolation compared to Baseline, Amygdala ROI.* Results are shown for a mixed effects regressions (using Matlab 2019b’s *fitlme* function) to estimate the fixed effects of cue, deprivation session, and their interaction on response magnitude in the ROI, controlling for each session’s average framewise displacement (i.e. head motion), with participant included as a random effect with both random intercepts and random slopes. The reported p-values (two-tailed) are not corrected for multiple comparisons.

##### Insula

###### Fasting vs Isolation contrast

| Predictors | Estimates | t | p |
| --- | --- | --- | --- |
| Session: Isolation > Fasting | 0.015 | 0.8 | 0.433 |
| Cue: Food | 0.165 | 7.1 | 1.47e-11 |
| Cue: Social | -0.223 | -9.6 | 9.81e-19 |
| Interaction: session * cue (food) | -0.034 | -2.0 | 0.052 |
| Interaction: session * cue (social) | -0.021 | -1.0 | 0.324 |

Supplementary Table 17. *Output mixed effects model: Fasting vs. Isolation contrast, Insula ROI.* Results are shown for a mixed effects regressions (using Matlab 2019b’s *fitlme* function) to estimate the fixed effects of cue, deprivation session, and their interaction on response magnitude in the ROI, controlling for each session’s average framewise displacement (i.e. head motion), with participant included as a random effect with both random intercepts and random slopes. The reported p-values (two-tailed) are not corrected for multiple comparisons.

###### Comparison to baseline session

| Predictors | Estimates | t | p |
| --- | --- | --- | --- |
| Session: Fasting | 0.020 | 0.6 | 0.583 |
| Session: Isolation | 0.051 | 1.1 | 0.266 |
| Cue: Food | 0.196 | 6.6 | 1.26e-10 |
| Cue: Social | -0.186 | -5.5 | 6.36e-8 |
| Interaction: session (fasting) – cue (food) | 0.004 | 0.1 | 0.924 |
| Interaction: session (isolation) – cue (food) | -0.065 | -1.8 | 0.070 |
| Interaction: session (fasting) – cue (social) | -0.016 | -0.4 | 0.701 |
| Interaction: session (isolation) – cue (social) | -0.057 | -1.4 | 0.151 |

Supplementary Table 18. *Output mixed effects model: Fasting and Isolation compared to Baseline, Insula ROI.* Results are shown for a mixed effects regressions (using Matlab 2019b’s *fitlme* function) to estimate the fixed effects of cue, deprivation session, and their interaction on response magnitude in the ROI, controlling for each session’s average framewise displacement (i.e. head motion), with participant included as a random effect with both random intercepts and random slopes. The reported p-values (two-tailed) are not corrected for multiple comparisons.

##### Anterior Cingulate Cortex (ACC)

###### Fasting vs Isolation contrast

| Predictors | Estimates | t | p |
| --- | --- | --- | --- |
| Session: Isolation > Fasting | 0.007 | 0.3 | 0.766 |
| Cue: Food | 0.117 | 3.6 | 0.0004 |
| Cue: Social | 0.034 | 1.0 | 0.337 |
| Interaction: session * cue (food) | -0.073 | -2.9 | 0.005 |
| Interaction: session * cue (social) | -0.007 | -0.3 | 0.787 |

Supplementary Table 19. *Output mixed effects model: Fasting vs. Isolation contrast, ACC ROI.* Results are shown for a mixed effects regressions (using Matlab 2019b’s *fitlme* function) to estimate the fixed effects of cue, deprivation session, and their interaction on response magnitude in the ROI, controlling for each session’s average framewise displacement (i.e. head motion), with participant included as a random effect with both random intercepts and random slopes. The reported p-values (two-tailed) are not corrected for multiple comparisons.

###### Comparison to baseline session

| Predictors | Estimates | t | p |
| --- | --- | --- | --- |
| Session: Fasting | 0.034 | 0.6 | 0.542 |
| Session: Isolation | 0.057 | 1.1 | 0.253 |
| Cue: Food | 0.139 | 3.4 | 0.0008 |
| Cue: Social | 0.022 | 0.5 | 0.634 |
| Interaction: session (fasting) – cue (food) | 0.052 | 1.1 | 0.273 |
| Interaction: session (isolation) – cue (food) | -0.095 | -1.8 | 0.078 |
| Interaction: session (fasting) – cue (social) | 0.019 | 0.3 | 0.736 |
| Interaction: session (isolation) – cue (social) | 0.005 | 0.1 | 0.908 |

Supplementary Table 20. *Output mixed effects model: Fasting and Isolation compared to Baseline, ACC ROI.* Results are shown for a mixed effects regressions (using Matlab 2019b’s *fitlme* function) to estimate the fixed effects of cue, deprivation session, and their interaction on response magnitude in the ROI, controlling for each session’s average framewise displacement (i.e. head motion), with participant included as a random effect with both random intercepts and random slopes. The reported p-values (two-tailed) are not corrected for multiple comparisons.

##### Group level whole brain: session * cue interaction

| Area | MNI coordinates  cluster peak  x y z | t value |
| --- | --- | --- |
| Right cerebellum | 14 -48 -46 | 5.90 |
| Left anterior cingulate cortex | -4 6 32 | 5.21 |
| Right occipital cortex | 26 -84 20 | 5.11 |
| Left cerebellum | -24 -68 -52 | 5.06 |
| Left superior parietal cortex | -26 -82 48 | 4.74 |
| Left midbrain (periaqueductal gray) | -8 -30 -12 | 4.73 |
| Left amygdala | -18 -2 -10 | 4.72 |
| Right premotor cortex | 4 -2 72 | 4.60 |
| Left dorsolateral prefrontal cortex | -30 38 30 | 4.57 |
| Left superior parietal cortex | -14 -54 62 | 4.52 |
| Right nucleus accumbens | 14 12 -10 | 4.46 |
| Right superior parietal cortex | 20 -54 60 | 4.06 |

Supplementary Table 21. *Group level whole brain session * cue interaction - Food: Fasting > Isolation.* Flexible factorial model using the first-level contrasts food > control and social > control from each session. Statistical inference was performed using a threshold of p < 0.05 corrected for multiple comparisons over the whole brain, using cluster-level correction. Supplementary Figure 4 (left) depicts the results from this table.

| Area | MNI coordinates  cluster peak  x y z | t value |
| --- | --- | --- |
| Left dorsomedial prefrontal cortex | -10 64 24 | 5.93 |
| Right occipital cortex | 34 -96 6 | 5.15 |
| Left caudate nucleus | -8 22 0 | 4.58 |
| Left orbitofrontal cortex | -2 40 -24 | 4.47 |
| Left occipital cortex | -14 -100 -8 | 4.19 |
| Left occipital cortex | -38 -94 -6 | 4.12 |
| Left occipital cortex | -26 -104 0 | 3.85 |

Supplementary Table 22. *Group level whole brain session * cue interaction - Social: Isolation > Fasting.* Flexible factorial model using the first-level contrasts food > control and social > control from each session. Statistical inference was performed using a threshold of p < 0.05 corrected for multiple comparisons over the whole brain, using cluster-level correction. Supplementary Figure 4 (right) depicts the results from this table.

##### Group level whole brain: main of effects cue

| **Food > Control** | | |
| --- | --- | --- |
| Area | Peak MNI coordinates  x y z | t value |
| Left fusiform gyrus | -30 -54 -12 | 19.04 |
| Left orbitofrontal cortex | -24 36 -14 | 15.01 |
| Left dorsolateral prefrontal cortex | -46 36 14 | 12.96 |
| Left anterior cingulate cortex | -4 4 30 | 11.68 |
| Right orbitofrontal cortex | 22 32 -16 | 10.37 |
| Right cerebellum | 18 -40 -44 | 10.29 |
| Left postcentral gyrus | -64 -18 30 | 9.98 |
| Left cerebellum | -22 -38 -42 | 9.81 |
| Left anterior cingulate cortex | -2 34 10 | 9.13 |
| Right parietal cortex | 28 -76 48 | 7.87 |
| Left perirhinal cortex | -24 -2 -32 | 7.74 |
| Left frontal cortex | -20 34 48 | 7.15 |
| Right fusiform gyrus | 52 -50 -18 | 6.19 |
| Right supramarginal gyrus | 66 -16 24 | 6.04 |
| Left frontal cortex | -8 50 52 | 5.47 |
| Left premotor cortex | -24 22 68 | 5.46 |
| Left perirhinal cortex | -18 -20 -24 | 5.30 |
| Right dorsolateral prefrontal cortex | 42 42 8 | 5.27 |
| Left premotor cortex | -20 24 70 | 5.23 |
| Left premotor cortex | -32 16 66 | 5.19 |
| Right parietal cortex | 26 -62 62 | 5.18 |
| Left caudate nucleus | -10 14 2 | 5.17 |
| **Social > Control** | | |
| Area | Peak MNI coordinates  x y z | t value |
| Left fusiform gyrus | 40 -48 -18 | 34.95 |
| Right inferior frontal gyrus | 48 22 22 | 12.77 |
| Right frontal cortex | 24 34 50 | 7.98 |
| Right premotor cortex | 48 2 58 | 7.46 |
| Right premotor cortex | 40 0 50 | 7.04 |
| Left posterior cingulate cortex | -4 -16 38 | 6.27 |
| Cerebellum | 0 -96 -24 | 5.78 |
| Left superior parietal cortex | -2 -64 -50 | 5.77 |
| Left cerebellum | -30 -64 -50 | 5.75 |
| Left frontal cortex | -4 36 64 | 5.55 |
| Left inferior temporal gyrus | -42 4 -44 | 5.50 |
| Left premotor cortex | -10 6 78 | 5.25 |
| Left premotor cortex | -14 28 68 | 5.24 |
| Right dorsolateral prefrontal cortex | 12 56 48 | 5.18 |

Supplementary Table 23. *Main effects of cue (food > control and social > control).* Flexible factorial model using the first-level contrasts food > control and social > control. Statistical inference was performed using a threshold of p < 0.05 corrected for multiple comparisons over the whole brain, using family-wise error correction at the voxel level. Supplementary Figure 6 depicts the results for this analysis.

1. Please note that the sign of the activation depends on the implicit baseline when modeling the data which makes the absolute value difficult to interpret (for a discussion on this issue, see Stark & Squire 2001^1^). When compared to the control condition (flower images), we see the expected higher activation in response to the craving cues (food and social).

   1	Stark, C. E. L. & Squire, L. R. When zero is not zero: The problem of ambiguous baseline conditions in fMRI. *Proceedings of the National Academy of Sciences* **98**, 12760-12766, doi:10.1073/pnas.221462998 (2001).
